# Msp1/ATAD1 restores mitochondrial function in Zellweger Spectrum Disease

**DOI:** 10.1101/2020.09.19.303826

**Authors:** Esther Nuebel, Jeffrey T Morgan, Sarah Fogarty, Jacob M Winter, Sandra Lettlova, Jordan A Berg, Yu-Chan Chen, Chelsea U Kidwell, J Alan Maschek, Katie J Clowers, Catherine Argyriou, Lingxiao Chen, Ilka Wittig, James E Cox, Minna Roh-Johnson, Nancy Braverman, Steven J Steinberg, Steven P Gygi, Jared Rutter

## Abstract

Peroxisomal Biogenesis Disorders (PBDs) are a class of inherited metabolic disorders with profound neurological and other phenotypes. The most severe PBDs are caused by mutations in peroxin genes, which result in nonfunctional peroxisomes typically through impaired protein import. In order to better understand the molecular causes of Zellweger Spectrum Disease (ZSD) -the most severe PBDs -, we investigated the fate of peroxisomal mRNAs and proteins in ZSD model systems. We found that loss of peroxisomal import has no effect on peroxin mRNA expression or translational efficiency. Instead, peroxin proteins—still produced at high levels— aberrantly accumulate on the mitochondrial membrane, impairing respiration and ATP generation. Finally, we rescued mitochondrial function in fibroblasts derived from human patients with ZSD by overexpressing ATAD1, an AAA-ATPase that functions in mitochondrial quality control. These findings might provide a new focus of PBD therapies in supporting quality control pathways that protect mitochondrial function.

## Introduction

Peroxisomes are single-membrane organelles that play an integral role in fatty acid metabolism, bile acid synthesis, and scavenging of reactive oxygen species [1]. For instance, plasmalogens, a class of lipids that act as precursors in myelin sheath formation, are solely synthesized in peroxisomes [2]. Peroxins are a group of membrane proteins that regulate peroxisomal biogenesis, inheritance, and metabolic processes. In addition to peroxisomal membrane proteins, more than 50 different enzymes are imported into the matrix of peroxisomes, where the majority of peroxisomal reactions occur. Given the multitude of metabolic pathways that require peroxisomes, it comes as no surprise that mutations in the majority of peroxin-encoding genes are linked to Peroxisomal Biogenesis Disorders (PBDs) with varying degrees of severity [3]. Patient cells can either have non-functional peroxisomes, decreased peroxisome biogenesis or function or even lack the organelle altogether. Clinical manifestation ranges from mild to severe [4]. In its most severe form, Zellweger Spectrum Disease (ZSD) can lead to myriad clinical features including seizures, hepatomegaly, renal cysts, skeletal abnormalities, and impaired hearing and eyesight [5], [6] and early childhood death. Current treatments are mainly supportive and focus on symptom management [6], [7].

Although we now know that the genomic mutations linked with ZSDs are universally in peroxin genes, Zellweger Syndrome was first associated with gross mitochondrial abnormalities [8]. Mitochondrial dysfunction is now widely recognized as a phenotype associated with ZSDs, however, the assumption has been that this is due to defects in peroxisomes and peroxisomal metabolism. Peroxisomes and mitochondria are physically and functionally coupled, as mitochondria not only share and complement peroxisomal roles in lipid metabolism and reactive oxygen species defense, but also regulate peroxisome biogenesis in conjunction with the endoplasmic reticulum [9],[10]. It is unclear whether these mitochondrial defects directly contribute to PBD pathophysiology, or are a secondary result of nonfunctioning peroxisomes, particularly in the brain and retina, which have high demand for mitochondrial metabolism [9],[10].

In the absence of functional peroxisomal import, peroxin levels must be decreased and regulated at the level of mRNA expression, translation, and/or protein stability or peroxin proteins -lacking their native destination—will necessarily mislocalize. Indeed, we previously found that at least some peroxins become targeted to the mitochondria when certain protein-sorting pathways such as the GET system, are impaired [13]. Yeast Msp1 and its mammalian homologue ATAD1, which belong to the AAA^+^ ATPase protein family, facilitates the extraction and degradation of mislocalized tail-anchored proteins from mitochondria [14], [15], [16]. This suggests that Msp1/ATAD1 might broadly facilitate the regulation of peroxin localization when peroxisomes are absent. However, the fate of all peroxins, as well as their regulation within ZSD model systems, is currently unknown. We therefore set out to understand how loss of peroxisomes affects peroxin gene regulation and if peroxin mislocalization to the mitochondria is a general feature in PBDs.

## Results

### Peroxin gene expression is maintained in the absence of functional peroxisomes

Using the budding yeast *S. cerevisiae* as a model system, we generated strains lacking either the peroxisomal biogenesis factor *PEX3* or *PEX19*, which have been previously described to cause loss of peroxisomes [17]. As mentioned above, some peroxin proteins mislocalize to the mitochondria and accumulate at that site when the mitochondrial quality control factor Msp1 is disrupted [13]. To test the effect of Msp1 loss in a PBD background, we also examined an *msp1Δ* single-deletion strain as well as *pex3Δmsp1Δ* and *pex19Δmsp1Δ* double-deletion strains. We monitored the presence of peroxisomes in each of these strains by expressing fluorescent proteins fused to the peroxisomal targeting motif, comprised of the amino acids SKL, which were imported by and thereby labeled the punctate peroxisomes present in wild-type and *msp1Δ* cells (Fig.1A). Conversely, we did not detect punctate peroxisomes in *pex3Δ* and *pex19Δ* cells (similarly for *pex3Δmsp1Δ* and *pex19Δmsp1Δ* double-deletion strains) (Fig.1A). Instead, faint fluorescence was visible throughout the cytosol, which indicates an absence of intact peroxisomes capable of protein import (Fig.1A). To confirm there is no remaining peroxisomal metabolic function despite the loss of punctate peroxisomes, we plated the strains on oleate-containing media, as yeast require functional peroxisomes to metabolize this otherwise toxic lipid. The *pex3Δ, pex19Δ, pex3Δmsp1Δ*, and *pex19Δmsp1Δ* strains grew normally on glucose and glycerol (Fig.1B, upper panels), but displayed a clear growth defect relative to wild type and *msp1Δ* yeast on oleate (Fig.1B, lower panels). These results indicate impaired peroxisome-mediated metabolism and suggest the loss of functional peroxisomes in these mutant yeast strains, which supports their use as a model for dissecting the molecular consequences of peroxisome loss.

**Figure 1.**
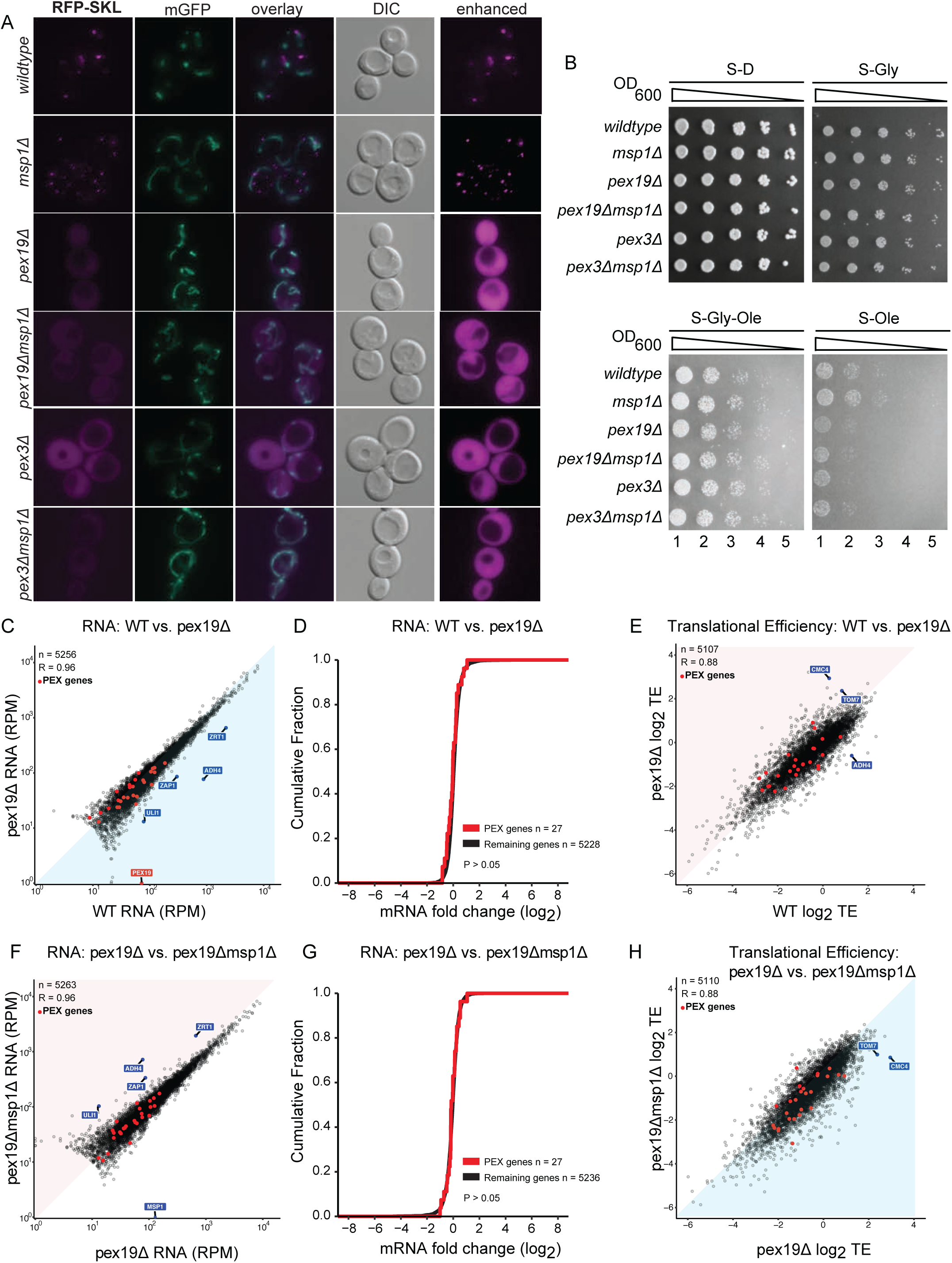
Peroxin gene expression is maintained in the absence of functional peroxisomes. (A) Yeast cells of each strain (*wild-type, msp1Δ, pex19Δ, pex19Δmsp1Δ, pex3Δ and pex19Δmsp1Δ*) expressing RFP-SKL (peroxisomal marker) construct and mitochondria-targeted green fluorescent protein (GFP) were grown to mid-log phase and analyzed by fluorescence microscopy. Representative images are shown. DIC, differential interference contrast. Enhanced: the red signal intensity was set to ‘best-fit’ in ZEN microscopy software analysis. (B) Yeast cells of each strain (*wild-type, msp1Δ, pex19Δ, pex19Δmsp1Δ, pex3Δ and pex19Δmsp1Δ)* were grown to mid-log phase, back diluted to 1 OD and serial dilutions were dropped onto agar plates containing synthetic media with dextrose (S-D), glycerol (S-Gly), glycerol and oleate (S-Gly-Ole) and oleate (S-Ole). (C) Wild-type versus *pex19Δ* RNA levels are blotted. Outliers are indicated. Red dots represent peroxin-RNAs. (D) Fold-change of RNA levels (log^2^) versus the cumulative fraction of wild-type versus *pex19Δ* is blotted. The red graph represents the peroxin distribution. (E) Translational efficiency (TE) of wild-type versus *pex19Δ*. Outliers are indicated. Red dots represent transaltional efficiency of peroxin genes. (F) *pex19Δ* versus *pex19Δmsp1Δ* RNA levels are blotted. Outliers are indicated. Red dots represent peroxin-RNAs. (G) Fold-change of RNA levels (log^2^) versus the cumulative fraction of *pex19Δ* versus *pex19Δmsp1Δ* is blotted. The red graph represents the peroxin distribution. (H) Translational efficiency (TE) of *pex19Δ* versus *pex19Δmsp1Δ*. Outliers are indicated. Red dots represent transaltional efficiency of peroxin genes.

We next sought to understand the regulation of peroxisomal genes in yeast that lack peroxisomes. It is unknown if an endogenous mechanism exists to downregulate expression of peroxisomal genes in the absence of functional peroxisomes. In addition to mRNA levels, we also considered the possibility of regulation at the level of translational efficiency (TE). Translation of some nuclear-encoded mitochondrial proteins is coupled to their import into the organelle [18], suggesting that imported peroxisomal proteins could similarly be regulated. Therefore, we performed RNA-Seq and ribosome-footprint profiling to measure differences in both gene expression and TE between wild-type and peroxisome-deficient *pex19Δ* yeast [19]. Despite lacking the highly conserved peroxisome, gene expression was remarkably similar in wild-type and *pex19Δ* yeast (Fig.1C). In particular, there was no significant difference in mRNA steady-state levels of peroxin-encoding genes (Fig.1D). A similar pattern was observed for TE as well (Fig.1E). Deletion of *MSP1* alone or in the *pex19Δ* background had no apparent effect on peroxin gene expression or TE, consistent with the known post-translational role of Msp1 in protein homeostasis (Fig.1F–H). Among the limited mRNA changes, we observed that three zinc-response genes (*ADH4, ZAP1*, and *ZRT1*) were downregulated in *pex19Δ* yeast relative to wild-type and *msp1Δpex19Δ* yeast (S-Fig.1A–D) [20]. This suggests a possible interplay between peroxisomal function and zinc homeostasis. Nonetheless, these results clearly demonstrate that the absence of peroxisomes does not significantly impact the steady-state expression or translation of peroxin-encoding genes.

### Peroxisomal proteins accumulate on mitochondria in the absence of peroxisomes

We next examined the abundance and localization of peroxin proteins under conditions where functional peroxisomes are absent using the *pex19Δ and pex3Δ* mutant strains. RNA-Seq and ribosome-footprint profiling data showed no substantial differences in peroxin transcription and translation in yeast lacking peroxisomes (in *pex3Δ* and *pex19Δ* backgrounds). Therefore, we hypothesize that either peroxin proteins are degraded more rapidly in these genetic backgrounds compared to wild-type or else peroxin proteins are mislocalizing *en masse* due to the absence of their expected destination. In the absence of protein sorting and mitochondrial quality control factors such as the gated entry of tail anchored proteins (GET) system and Msp1, some peroxins can become mislocalized to the mitochondrial outer membrane, suggesting it may also serve as a cellular destination for peroxins in the absence of peroxisomes [13]. To explore this hypothesis, we fused two representative peroxins, Pex11 and Pex13, to RFP and determined their localization via microscopy. Pex11-RFP and Pex13-RFP both co-localize with peroxisome markers in wild-type yeast, but co-localize with a mitochondrial marker in the *pex19Δ*, which colocalization becomes more dramatic in the *pex19Δmsp1Δ* double mutant strain (Fig.2A,B). These data suggests that peroxins can accumulate on the mitochondrial surface in cells lacking functional peroxisomes.

**Figure 2.**
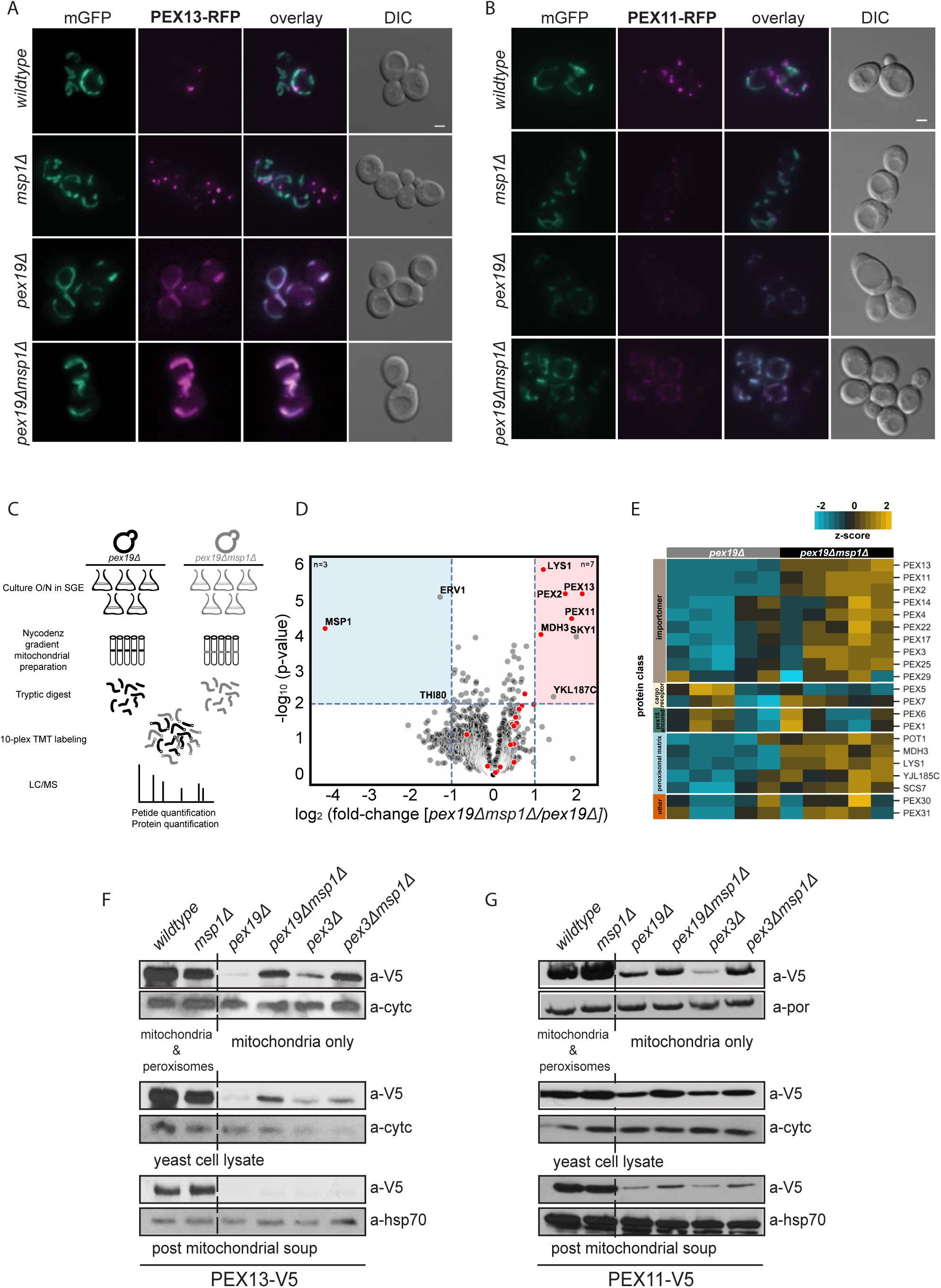
Peroxisomal proteins accumulate on mitochondria in absence of peroxisomes in yeast. (A) Yeast cells of each strain expressing the PEX13-RFP construct and mitochondria-targeted green fluorescent protein (GFP) were grown to mid-log phase and analyzed by fluorescence microscopy. Representative images are shown. DIC, differential interference contrast. (B) Yeast cells of each strain expressing the PEX11-RFP construct and mitochondria-targeted green fluorescent protein (GFP) were grown to mid-log phase and analyzed by fluorescence microscopy. Representative images are shown. DIC, differential interference contrast. (C) Depiction of the sample generation for quantitative mass spectrometry (experimental flow). (D) Volcano plot representing the average of 5 biological replicates of each strain (pex19Δ and pex19Δmsp1Δ) to indicate most enriched/ decreased proteins in the mitochondrial proteome of pex19Δ and pex19Δmsp1Δ. Detected by quantitative mass spectrometry. P-values were corrected using the Benjamini-Hochberg adjustment procedure. Gray dots represent all measured proteins, and red dots highlight peroxisomal-associated proteins. (E) Heatmap representing 5 biological replicates of each strain (pex19Δ and pex19Δmsp1Δ) and their peroxin proteinlevels detected by quantitative mass spectrometry, protein classes are indicated in the row labels. Values were z-score normalized by protein. (F) Total yeast cell lysate, post mitochondrial soup and Histodenz^™^ purified mitochondria of each strain (pex19Δ and pex19Δmsp1Δ; pex3Δ pex3Δmsp1Δ) transformed with Pex13^V5^ (expressed by its endogenous promotor) were separated by SDS-PAGE and immunoblotted for Pex13 (a-V5V5) and porin (a-porin), cytochrome c (cyt c) and HSP70 (hsp70). (G) Total yeast cell lysate, post mitochondrial soup and Histodenz^™^ purified mitochondria of each strain (pex19 and pex19msp1; pex3 pex3msp1) transformed with Pex11^V5^ (expressed by its endogenous promotor) were separated by SDS-PAGE and immunoblotted for Pex11 (a-V5) and porin (a-porin), cytochrome c (cyt c) and HSP70 (hsp70).

To extend this finding and better understand the fate of peroxin proteins in general, we next performed quantitative mass spectrometry of mitochondria isolated by Histodenz^™^ gradient purification from yeast lacking peroxisomes. Isobaric labeling of tryptic peptides then allowed for a quantitative measure of the abundance of each detected protein (Fig.2C) [21]. Peroxisomes and mitochondria are known to form extensive contact sites [22], and isolated mitochondria from wild type and *msp1Δ* strains would be contaminated with peroxisomes. Therefore, we chose to focus our studies of the mitochondrial proteomes of only peroxisome-lacking strains (*pex19Δ* and *pex19Δmsp1Δ*). Therefore, this experiment can address whether loss of the mitochondrial quality-control protein Msp1 results in an increase in peroxins associated with mitochondria. Indeed, we found increased abundance of 12 peroxins, as well as 5 peroxisomal matrix proteins in the mitochondrial proteome of *pex19Δmsp1Δ* yeast compared to *pex19Δ* yeast (Fig.2D,E). To exclude that this result is *pex19Δ*-specific, we performed the same experiment comparing the mitochondrial proteome of *pex3Δ* to *pex3Δmsp1Δ* yeasts with similar results (S-Fig.2A,B).

To validate the quantitative mass spectrometry results, we V5-tagged two highly enriched peroxins (Pex11 and Pex13), transformed the constructs into wild-type, *msp1Δ, pex19Δ*, and *pex19Δmsp1Δ* strains, isolated intact mitochondria by Histodenz^™^ gradient, and performed immunoblotting. As expected, strains containing mitochondria and peroxisomes (wild type and *msp1Δ*) showed a strong signal for the V5-tagged proteins (Fig.2F,G), which is due to peroxisomal contamination caused by co-purification of both organelles [22]. The *pex19Δ* mitochondria displayed a barely detectable signal for both Pex13-V5 and Pex11-V5; comparatively, both Pex13-V5 and Pex11-V5 were dramatically enriched in *pex19Δmsp1Δ* mitochondria (Fig.2F,G). Again, similar results were obtained with *pex3Δ* and *pex3Δmsp1Δ* mitochondria (Fig.2F,G). We performed an identical experiment with Pex30-V5—a peroxin that was shown to accumulate on the ER—and found that Msp1 had no effect on causing Pex30 to accumulate on mitochondria instead (S-Fig.2C). This confirms the specificity of peroxins detected on mitochondria. In all, our mass spectrometry and Western blotting data show that many peroxisomal proteins accumulate on mitochondria in the absence of peroxisomes, and that this accumulation is exacerbated by loss of the mitochondrial quality control protein Msp1.

### Pex13 mediates docking and accumulation of peroxin complexes on mitochondria

Peroxin proteins assemble into specific complexes in the peroxisomal membrane, which enable protein import into the peroxisomal matrix through known docking and import pathways. Pex13 is part of the docking subcomplex of the peroxisomal importomer. Because Pex13 is mislocalized to mitochondria in peroxisome-deficient strains lacking Msp1, we reasoned that it might serve to dock peroxisomal proteins to mitochondria as it normally would to peroxisomes [21], [22]. Therefore, we asked whether peroxins that become localized to mitochondria still assemble into native complexes, and whether those peroxins required for this process on peroxisomes are similarly required for the recruitment of peroxins to mitochondria. To address this question, we first isolated mitochondria from yeast strains expressing Pex13-V5—which accumulates on the mitochondria in peroxisome-deficient strains (Fig.2)—and separated them on a 3-18% gradient Blue Native PAGE using different amounts of detergent (Fig.3A,B). To detect Pex13-containing complexes, we titered the amount of digitonin detergent in our mitochondrial purification protocol in order to optimize the protection of native protein complexes while effectively disrupting non-specific interactions, which we monitored using internal controls [23](Fig.3B,C). At four grams of digitonin per gram of mitochondria, Pex13-V5 migrates in a complex with a total mass of ∼180 kD (Fig.3C,D). This mass is consistent with that of the native docking complex (172.5 kD), which is made up of Pex13, Pex14, Pex17, and Pex8. As expected, this complex is abundant in wild-type yeast (Fig.3C, black box) with intact peroxisomes and is absent in peroxisome-deficient *pex19Δ* yeast. Interestingly, this complex is restored upon additional deletion of *MSP1* in *pex19Δmsp1Δ* yeast (Fig.3C, grey box). These data indicate that at least one native peroxin complex forms in the absence of peroxisomes on mitochondria (Fig.3E).

**Figure 3.**
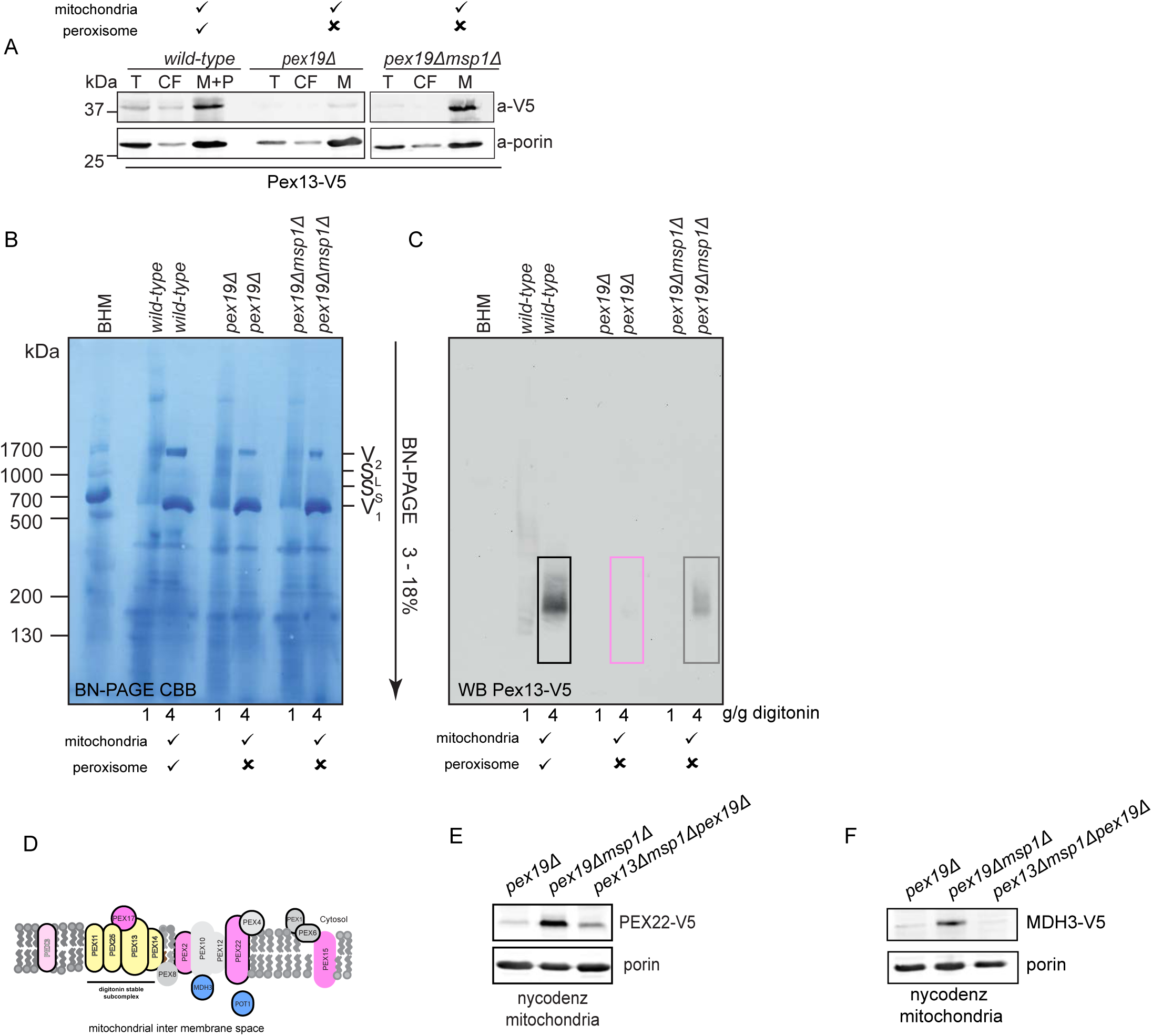
Pex13 mediates docking and accumulation of peroxin complexes on mitochondria. (A) Fractionated yeast cells (total = T, post mitochondrial soup = PMS, mitochondria = mito) expressing PEX13^V5^ (endogenous promotor) from yeast strains (wildtype, pex19Δ, pex19Δmsp1Δ) were separated by SDS-PAGE and immunoblotted for PEX13^V5^ (V5) and Porin (porin). (B) Digitonin solubilized (1 and 4 g/g detergent/ membrane ratio as indicated) mitochondrial membranes expressing PEX13^V5^ (endogenous promotor) from yeast strains (wildtype, pex19Δ, pex19Δmsp1Δ) were separated by BN-PAGE with a 3-18% gradient stained with Coomassie brilliant blue. (C) Digitonin solubilized (1 and 4 g/g detergent/ membrane ratio as indicated) mitochondrial membranes expressing PEX13^V5^ (endogenous promotor) from yeast strains (wildtype, pex19Δ, pex19Δmsp1Δ) were separated by BN-PAGE with a 3-18% gradient. Gel was transferred to PVDF membrane and immunoblotted for PEX13^V5^ (V5). (D) Depiction of detected peroxins which contribute to the peroxisomal importomer assembled on mitochondria. Cartoon representation. (E) Total yeast cell lysate, post mitochondrial soup and Histodenz^™^ purified mitochondria of each strain (pex19Δ and pex19Δmsp1Δ and pex13Δpex19Δmsp1Δ) transformed with Pex22^V5^ (expressed by its endogenous promotor) were separated by SDS-PAGE and immunoblotted for Pex22 (a-V5) and porin (a-porin) (F) Total yeast cell lysate, post mitochondrial soup and Histodenz^™^ purified mitochondria of each strain (pex19Δ and pex19Δmsp1Δ and pex13Δpex19Δmsp1Δ) transformed with Mdh3^V5^ (expressed by its endogenous promotor) were separated by SDS-PAGE and immunoblotted for Mdh3 (a-V5) and porin (a-porin)

With this data suggesting that a native peroxisomal docking complex forms on mitochondria, we next wanted to know whether this complex is functional and contributes to the recruitment and mitochondrial localization of additional peroxins. To address if Pex13 is required for recruitment of other peroxins to the mitochondria, we created *pex13Δ* strains in the background of *pex19Δ, msp1Δ*, and *pex19Δmsp1Δ* (S-Fig.3). To track peroxin localization in these strains, we V5-tagged two additional peroxisomal proteins: Pex22 and Mdh3. We found that both Pex22-V5 and Mdh3-V5 co-fractionate with mitochondria in a *pex19Δmsp1Δ* mutant, similar to the pattern observed for Pex 13 and Pex11 previously. This localization to mitochondria was reduced to background levels when Pex13 was removed (Fig.3F,G). It is particularly intriguing that we see Pex13-dependent and Msp1-sensitive localization of Mdh3 to mitochondria as Mdh3 is a soluble, peroxisomal matrix protein. Together, these results indicate that, in the absence of peroxisomes, a functional peroxisomal docking and import complex forms on mitochondria.

Further, the docking protein Pex13—and possibly other components of these complexes—is required for the recruitment of additional peroxisomal proteins. Since the overwhelming accumulation of peroxins on the mitochondria is likely responsible for mitochondria dysfunction in ZSDs [24],[25], we hypothesize that abolishment of the docking complex formation particularly on mitochondria will be beneficial for mitochondrial function.

### Peroxins target to mitochondria in cells derived from Zellweger Spectrum Disorder patients

Thus far, we have used yeast as a model system to study ZSDs. Although peroxisome function and peroxin proteins are highly conserved among eukaryotes, we wanted to test whether our findings in yeast are representative of the human ZSD scenario. To do this, we started with a PEX3 deficient (herein referred to as PEX3^-/-^) fibroblast cell line derived from a patient with Zellweger Syndrome. To create an isogenic control cell line, we complemented this cell line by expressing wild-type PEX3 (cWT) (S-Fig.4A). In line with published data, this restores peroxisome formation and function [10]. We then overexpressed ATAD1—the human ortholog of yeast Msp1—to create a total of four cell lines: cWT, cWT+ATAD1, PEX3^-/-^, PEX3^-/-^ +ATAD1. Finally, in order to track peroxin localization, we expressed PEX13-GFP using a lentiviral vector in these four cell lines.

In cWT, and cWT + ATAD1 cells, which should both contain functional peroxisomes, PEX13-GFP displayed a bright, punctate pattern, in agreement with its expected peroxisomal localization (Fig.4A). In PEX3^-/-^ cells, the PEX13-GFP signal was slightly dimmer but entirely overlapping with the mitochondrial network, which indicates peroxins are similarly localized to mitochondria in human cells derived from ZSD patients. In PEX3^-/-^+ATAD1 cells, GFP was no longer found at mitochondria, but was instead found throughout the cytosol and nucleus—a pattern also found in control cells expressing free GFP (S-Fig.4A lanes 1 to 4) and probably explained by the observation that the PEX3^-/-^+ATAD1 cells exhibit primarily free GFP (S-Fig.4A lane 8). We performed an analogous set of experiments in a PEX16-deficient cell line derived from a second Zellweger Syndrome patient, in this case we transiently expressed PEX10-GFP and PEX13-GFP to track peroxin localization (S-Fig.4B,C) [26] and found colocalization of both peroxins in the mutant fibroblast with mitochondria. The results were similar to those reported here, and therefore support the biology of peroxin localization to mitochondria a broad array of ZSDs.

These experiments confirm that, as shown in yeast previously, peroxins also localize to mitochondria in human cells lacking functional peroxisomes—in this case, cells derived from patients with Zellweger Spectrum Disease. Additionally, they show that overexpression of ATAD1 is able to ameliorate much of the peroxin mislocalization to the mitochondria, akin to its function in removing tail anchored proteins and other substrates by extracting them from the outer mitochondrial membrane [13], [27].

### Overexpression of ATAD1 rescues mitochondrial dysfunction in human ZSD fibroblasts

Based on our findings that peroxins localize to mitochondria in cells derived from Zellweger Syndrome patients, and that overexpression of ATAD1 results in removal of peroxins from mitochondria, we next asked whether ATAD1 overexpression is able to improve mitochondrial function in these cells. We started by classifying gross mitochondrial morphology using electron microscopy. Cristae are barely detectable in PEX3^-/-^ mitochondria; in addition, based on osmium staining, mitochondria are more protein dense compared to wild-type controls, consistent with aberrant protein accumulation at mitochondria (Fig.4B) [28]. To quantify mitochondrial morphology, we outlined three categories: mitochondria with many cristae and low electron density (type 1), mitochondria with fewer cristae and low electron density (type 2) and mitochondria with few or no cristae and high electron density (type 3-damaged mitochondria). Whereas 80% of mitochondria in wild-type cells match the criteria for types 1 and 2, more than 80% of mitochondria in PEX3^-/-^ patient cells instead were quantified as type 3. Overexpression of ATAD1 in the PEX3^-/-^ background largely rescued the morphological defects, however, as 70% of mitochondria matched the criteria for type 2 (S-Fig.4D). Thus, based on gross mitochondrial morphology, overexpression of ATAD1 was sufficient to restore normal mitochondria morphology in cells derived from Zellweger Syndrome patients.

Given that overexpressing ATAD1 normalized mitochondrial morphology in peroxisome-deficient cells, we next set out to determine if mitochondrial metabolic function might also be restored. First, we measured respiration using a Seahorse XF Flux Analyzer (Fig.4C–F). PEX3^-/-^ cells had both a lower basal rate of respiration and a lower maximal respiratory capacity compared to wild-type cells (Fig.4C–F), which agrees with their damaged appearance by EM (Fig.4B). ATAD1 overexpression in PEX3^-/-^ cells was able to restore both basal mitochondrial respiration and maximal respiratory capacity to wild-type levels (Fig.4C–F). Additionally, we generated an *ATAD1* knockout cell line in the PEX3^-/-^ background, which exhibited even further reduced mitochondrial respiration compared to its parental PEX3^-/-^ control (Fig.4C–F). These results indicate that even endogenous ATAD1 is acting to maintain mitochondrial function in PBDs to an extent (as observed in the PEX3^-/-^ cell line). It is seemingly not sufficient, however, to combat the overwhelming amount of peroxins that become mislocalized to mitochondria. Therefore, overexpression of ATAD1 is required to restore mitochondrial respiration and morphology to wild-type levels in these cells (PEX3^-/-^+ ATAD1).

Lastly, we aimed to investigate features related to mitochondria and metabolism other than respiration and performed whole-cell lipidomics (S-Table1). Specific lipid classes, such as cardiolipins and phosphatidylethanolamines (PE), are associated with active mitochondrial metabolism. Indeed, we found cardiolipins, which are depleted in PEX3^-/-^ cells, are restored to wild-type levels by overexpression of ATAD1 and phosphatidylethanolamines exhibit an overall increase (Fig.4G,H). In addition, we found that 30% of quantified phosphatidylethanolamines are lower in the patient cell line, but all of those PE species increase with ATAD1 expression (Fig.4I). Plasmalogens, on the other hand —which are precursors in myelin synthesis—are similarly decreased in PEX3^-/-^ cells but cannot be rescued by ATAD1 overexpression. This comes with no surprise since their synthesis requires functional peroxisomes (Fig.4H). These results provide further evidence that overexpression of ATAD1 specifically rescues many aspects of mitochondrial dysfunction in human PBD fibroblasts.

## Discussion

It has been well described in the literature that a diverse array of eukaryotic cells and organisms can survive without functional peroxisomes [29]. Yet, human PBDs present with severe and rapidly progressive phenotypes that greatly limit patients’ quality of life [6]. Our study therefore sought to understand the molecular underpinnings of PBDs’ cellular phenotypes, focusing on the regulation and fate of peroxisomal mRNAs and proteins when peroxisomes are absent. Previous studies in yeast have found that transcription and translation of mitochondria-destined proteins are decreased when mitochondrial import pathways are dysfunctional [30], raising the possibility that a similar regulatory paradigm could be present for peroxins. However, RNA-Seq and ribosome-footprint profiling showed that peroxin gene expression as well as peroxin mRNA translational efficiency are unaffected by stable, genetic loss of peroxisomes. Of note: a previous study reported redistribution of peroxin mRNA from the cytosol to the periphery of peroxisomes when cells are shifted to media that induces rapid peroxisomal biogenesis [31]. So, although we do not see changes in translational efficiency upon loss of peroxisomes, this does not exclude the possibility that peroxins as a class of mRNA might demonstrate coordinated regulation of translational efficiency upon changes to the cellular environment and/or metabolic needs.

Using both microscopy and high-throughput proteomics, we found that many peroxins— most prominently Pex13, Pex11, Pex14, Pex2, Pex17 and Pex25—accumulate on the mitochondrial outer membrane when peroxisomes are absent. This accumulation led to the formation of the peroxisomal docking complex, a subcomplex of the peroxisomal import machinery [32],[33],[34],[35], on mitochondria. The formation of this complex was shown to play a pivotal role in recruiting both peroxisomal membrane proteins and peroxisomal matrix proteins, such as Mdh3, to the mitochondria. To our knowledge this is the first evidence of a functional membrane protein translocase localizing to another organelle and thereby putatively interfering with its function. Further work will be needed to uncover if the peroxisomal import machinery has a distinct array of substrates when localized to the mitochondria *versus* the peroxisome.

In the field of peroxisomal biology, two main mechanisms of peroxisomal biogenesis are widely accepted: the multiplication of peroxisomes initialized by a growth and division pathway as well as *de novo* peroxisome generation involving the endoplasmic reticulum (ER) [36][37][38][39]. A third mechanism has been postulated in which mitochondrial derived vesicles contribute to peroxisome biogenesis [10]. Our findings of peroxisomal proteins on mitochondria, albeit in a disease background, might provide new perspectives into the ways in which peroxisomes are formed, their substrates are imported and mitochondrial quality control mechanisms regulate these processes. While we focused this work on a disease setting, it is worth further exploration of these processes within the context of a healthy cell environment.

Although peroxins represented the clearest hits from our proteomics data, some other proteins are worthy of further discussion. Particularly, we found three zinc-related proteins (Zrt1, Zap1, Adh4) that were reciprocally regulated in *pex19Δ* compared to wild-type *pex19Δmsp1Δ* yeast. Zrt1 is a zinc transporter, Zap1 is a zinc-sensitive transcription factor and zinc sensor, and Adh4 is a zinc-sensitive alcohol dehydrogenase. In addition to these three proteins, we also saw that known Zap1 target genes (including ADH4) were similarly more abundant at the RNA level in wild-type yeast than in *pex19Δ* yeast (Fig.1E and S-Fig.1E). Although we are not aware of a well-characterized connection between peroxisome function and Zn^2+^ content, several peroxins contain zinc-binding domains, and mutations to these domains in Pex2, Pex10, and Pex12 cause particularly severe peroxisomal defects and PBD phenotypes [40],[41],[42]. Therefore, it is possible that there is a regulatory connection between Zn^2+^ homeostasis and peroxin function.

Despite loss of a highly conserved organelle, gene expression and translational efficiency measurements were remarkably similar between wild-type and *pex19Δ* yeast. However, as with the proteomics data, there are a small number of mRNAs that display differential translational efficiency in these two strains. In particular, *CMC4* and *TOM7* appear to be translated more efficiently in *pex19Δ* yeast than in wild-type yeast (Fig.1E,H). The protein products of both of these genes are involved in or dependent on the Mia40/ERV1 mitochondrial import pathway [43], which we found to be affected in our *pex* mutant strains (Fig.2D). Why these two particular mitochondrial genes would be post-transcriptionally regulated in this way is unclear. This suggests to us that different mitochondrial import pathways could be affected by peroxin mislocalization and warrants further investigation.

We are particularly intrigued by the implications of our findings on Msp1/ATAD1. We show that overexpression of ATAD1 is sufficient to restore mitochondrial morphology, respiration, and metabolism in cells derived from ZSD patients. Our study suggests that therapies that can similarly increase ATAD1 expression, function (or provide similar benefits via distinct mechanisms) may relieve the burden of peroxins and restore function to mitochondria in ZSD patients. Insofar as Msp1/ATAD1 protects mitochondria in ZSD cells by dismantling the mislocalized peroxisomal import apparatus, a drug that interferes with this import process might achieve similar therapeutic benefits. In other words, it may be that the only peroxisomal import that occurs in ZSD cells is aberrant import into mitochondria; in that case, inhibiting peroxisomal import could actually benefit ZSD patients. A small-molecule inhibitor of trypanosomal PEX14 was recently described to kill T. brucei parasites by blocking peroxisomal protein import [44]. Perhaps repurposing similar compounds could be an unconventional pharmacological approach in PBD patients. Beyond the direct benefits to mitochondrial dysfunction at large, improvements to cells’ metabolic function might allow more aggressive, combinatorial therapies to help alleviate the plethora of other ZSD symptoms. Overall, this work suggests that augmenting mitochondrial quality control might be a promising therapeutic strategy for ZSDs.

## Methods

### Yeast Strains

*Saccharomyces cerevisiae* W303 (MATa/α, +/ade2 +/arg4 +/can1 +/lys2 +/met15 trp1-1/trp1delta0 ura3-1/ura3delta0), was used to generate all knockout strains. To generate haploid single and double mutants JRY2722 msp1::KanMX/MSP1 pex19::natMX/PEX19 (MATa/α, his3 / his3 leu2 / leu2 lys2 / lys2 met15 / met15 trp1 / trp1 ura3 / ura3) and JRY2723 msp1::KanMX/MSP1 pex19::natMX/PEX19 (MATa/α, his3 / his3 leu2 / leu2 lys2 / lys2 met15 / met15 trp1 / trp1 ura3 / ura3) were generated using a standard PCR-based homologous recombination method. Briefly, the drug selection cassette (KanMX4, hphMX4, or natMX4) flanked with ca. 45-bp fragments upstream and downstream of the gene of interest was PCR amplified and transformed into the wildtype diploid strain [45], [46]. JRY4461 was generated by crossing JRR4470 with JRY2726. Following, the haploid strains were generated by sporulation and tetrad analysis. The genotype of the strains was verified by standard genotyping PCR. The genotypes of all yeast strains used in this study are listed in the Key Resources Table. Yeast transformation was performed by the standard TE/LiAc method [47] and transformed cells were recovered and grown in synthetic complete glucose (SD) medium lacking the appropriate amino acid(s) for selection purposes. Medium used in this study is synthetic minimal medium supplemented with different carbon sources: 2% glucose, 2% glycerol, 2 % ethanol, 0.1% oleate and appropriate amino acids. Solid plates contain 2.2% (w/v) agar.

### Plasmid construction

To produce plasmids expressing non-tagged, C-terminal His6/HA3, -V5 or GFP/ RFP-tagged peroxins or Msp1, the ORF flanked with its upstream ca. 600-bp promoter region was PCR amplified from yeast genomic DNA and ligated into the pRS416, pRS426, or pRS416 vector containing either a C-terminal His6/HA3, V5, or GFP/ RFP tag. To generate human ATAD1 constructs, the ATAD1 ORF was amplified from HepG2 and ligated into pRS416-based vector containing the ADH1 promoter [48] as described in [13]. The Retro-XTMQ Vectors (Clontech) were used to generate mammalian constructs. Human ATAD1 ORF fused with HA or GFP-tag at the 30-end or PEX3, PEX13 or PEX11 ORF fused with GFP at the 50-end was ligated into pQCXIP or pQCXIZ vector. Peroxisomal fluorescent marker construct was made by fusing RFP protein with the peroxisomal targeting sequence (SKL) at the C-terminus.

### Yeast Growth Assays

Growth assays were performed using synthetic minimal media supplemented with the appropriate amino acids and indicated carbon source. For plate-based growth assays, overnight cultures were back-diluted to equivalent ODs and spotted as 5-fold serial dilutions. Cells were allowed to grow for 2-3 days at 30°C before imaging.

### Ribosome profiling

RNA-seq and ribosome-footprint profiling (RPF). Libraries were prepared as described in [49] with small modifications from [50]. Briefly, yeast cultures were grown to mid-log phase, collected by vacuum filtration, and rapidly flash-frozen in liquid nitrogen. Lysate powder was aliquoted and stored at −80 °C. Frozen pellets were mechanically lysed using a Sample Prep 6870 Freezer/Mill (Spex SamplePrep; 10 cycles of 2 min on, 2 min off at setting 10).

For RNA-seq samples, total RNA was extracted from lysate powder using TRI Reagent (Ambion) according to the manufacturer’s protocol. rRNA was depleted from 5 μg of total RNA using the Ribo-Zero Gold Yeast rRNA Removal Kit (Illumina) according to the manufacturer’s protocol. For RPF samples, lysate powder was resuspended in a buffer containing cycloheximide and tigecycline in order to arrest translating ribosomes. Monosomes were isolated by treating the lysate with RNase I followed by purification over a 10-50% sucrose gradient. Ribosome-protected fragments were further isolated by Proteinase K/SDS digestion and phenol/chloroform extraction.

RNA-seq samples were fragmented by alkaline hydrolysis and size-selected in the range of 19-40 nt. RPF samples were size-selected in the range of 19-33 nt, and contaminating rRNA sequences were removed via subtractive hybridization. RNA-seq and RPF samples were both ligated to adaptors, reverse-transcribed, and amplified to prepare sequencing libraries. These libraries were sequenced on an Illumina HiSeq 2500 using 50-bp single-end reads.

### Fluorescence microscopy

WT W303 (245) or derived mutant strains were transformed with a plasmid expressing mitochondria-targeted (ATPase subunit, Su9) GFP, mtGFP, and plasmids expressing Pex13-RFP, RFP-SKL, or Pex11-RFP under their endogenous promoter, respectively. Samples were prepared by growing yeast cells to early log phase (OD600 = 0.8–1) in synthetic dropout medium with appropriate carbon sources at 30°C. To prepare mammalian cell samples, we cultured cells on chamber slides for a day to allow them to fully attach. They were either visualized directly or stained with 25 nM Mitotracker Red CMXRos (Life Technologies) in FBS-free medium for 10 min, followed by a 10 min recovery in serum containing media which was ultimately changed to phenol red free media at 37°C before imaging. Transfected alive cells were imaged after staining with Mitotracker red. Green signals were acquired by tagging the transfected peroxin with GFP. Images of the cells were acquired by using a Zeiss LSM 710 laser-scanning confocal microscope using a 40 or 100Å∼ oil-immersion objective. Cells were imaged on the Axio Observer. Z1 imaging system (Carl Zeiss) equipped with 40Å∼ and 100Å∼ objectives (oil-immersion). Digital fluorescence and differential interference contrast (DIC) images were acquired using a monochrome digital camera (Axio-Cam MRm, Carl Zeiss). ZenPro software (Version 4.8, CarlZeiss) was used to optimize images (tool: best fit). The final images were assembled using Adobe Photoshop CS5.1. The yeast fluorescent images shown in this study are representative pictures from at least two independent experiments (n ≥ 2). The images of the mammalian cells are representative of at least two independent experiments (n ≥ 2).

### Airy Scan microscopy

The Zeiss LSM880 with Airyscan (Carl Zeiss, Germany) was used to acquire images in figure 4 A. Mammalian cells were plated overnight on glass bottom fluorodishes (WPI Inc), and imaged under 37C and 5% CO_2_ conditions. Z-stacks were acquired with a 63X 1.4 NA oil objective using Immersol 518F /30C immersion fluid (Carl Zeiss, Germany) under Fast Airyscan mode [46]. Raw images were Airyscan processed using Zen software (Carl Zeiss, Germany) with ‘auto’ settings. FIJI software [47] was then used to generate a maximum projection image comprising X μm of the sample, and linear adjustments were made to the brightness and contrast of each image.

**Figure 4.**
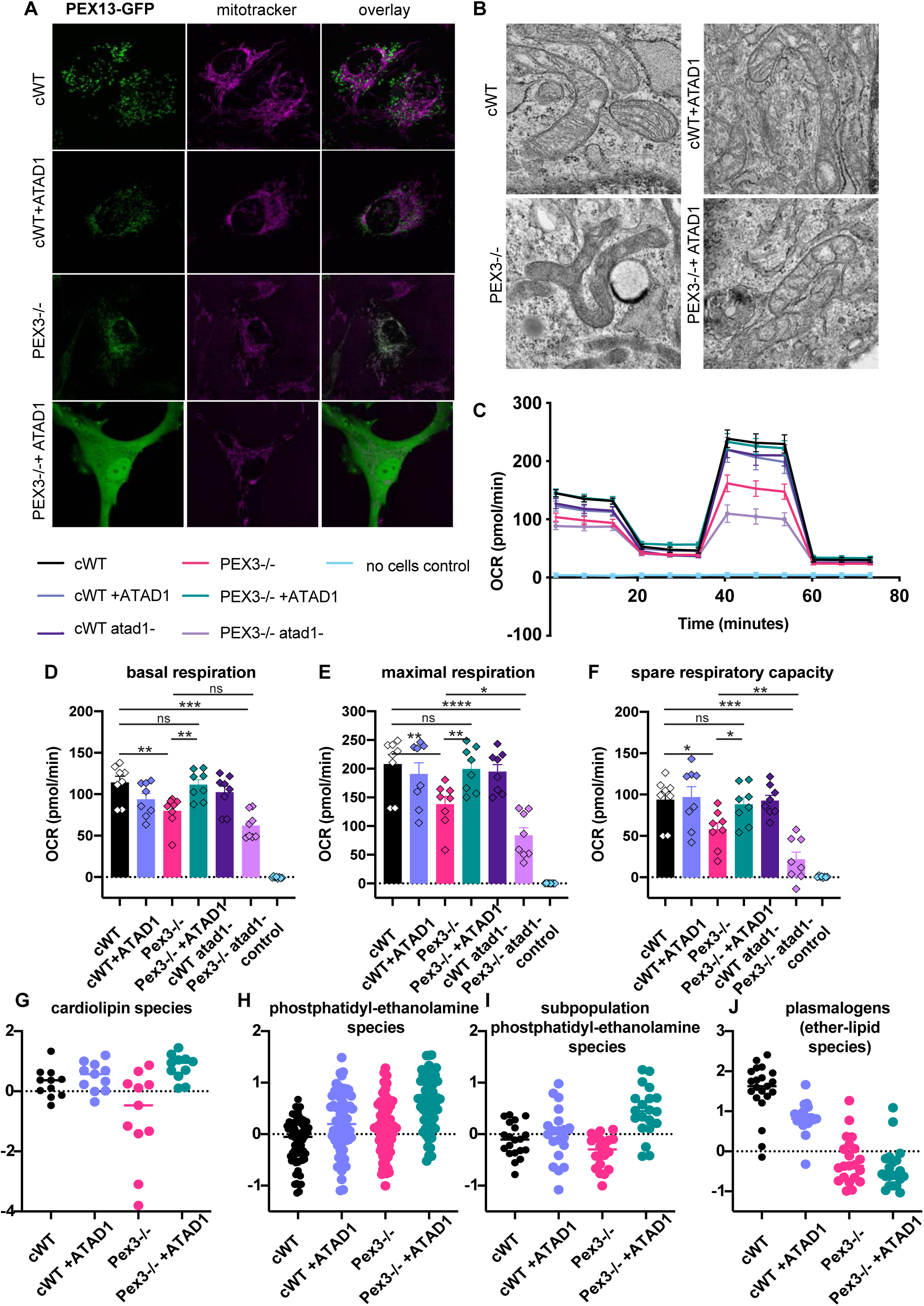
Peroxins target to mitochondria in cells derived from ZSD patients and overexpression of ATAD1 rescues their mitochondrial dysfunction. (A) Live fluorescence microscopy of patient fibroblast cell lines: cWT (wild-type + empty vector), cWT+ATAD1 (wild-type expressing ATAD1) PEX3-/-(patient + empty vector), PEX3-/-+ATAD1 (PEX3-/-expressing ATAD1) expressing PEX13-GFP stained with Mitotracker far red (MT) to visualize the mitochondrial network and GFP fused to PEX13 (PEX13-GFP) to investigate the localization. (B) Electron microscopy of patient fibroblast cell lines: cWT (wild-type + empty vector), cWT+ATAD1 (wild-type expressing ATAD1) PEX3-/-(patient + empty vector), PEX3-/-+ATAD1 (PEX3-/-expressing ATAD1). Representative images of the most observed mitochondrial morphology are shown. (C) Bioenergetic profile of human patient fibroblasts. OCR (pmol/min/norm.unit) for cWT (wild-type + empty vector), cWT+ATAD1 (wild-type expressing ATAD1) PEX3-/-(patient + empty vector), PEX3-/-+ATAD1 (PEX3-/-expressing ATAD1), cWT-ATAD1 (wildtype with ATAD1 deletion), PEX3-/-atad1 (patient with ATAD1 deletion) cells plotted against time (XF96e Extracellular Flux Analyzer, Mito-Stress-Test.) Oligomycin A (1 μM), 0.15 μM FCCP and 1 μM rotenone + 1 μM antimycin A (final concentrations) were sequentially delivered to the XF96e assay medium through injection ports in the sensor cartridge. The graph is the compilation of 2 independent assays (n=4+4=8 technical replicates). (C) Bar graph representation of basal respiration measured in (B) cWT (wild-type + empty vector), cWT+ATAD1 (wild-type expressing ATAD1) PEX3-/-(patient + empty vector), PEX3-/-+ATAD1 (PEX3-/-expressing ATAD1), cWT-ATAD1 (wildtype with ATAD1 deletion), PEX3-/-atad1-(patient with ATAD1 deletion). Statistical significance was calculated using the Welch’s test, p values are indicated as: ns (not significant), * p≤0.05, ** p≤0.01, *** p≤0.001, **** p≤0.0001. (D) Bar graph representation of maximal respiration measured in (B) cWT (wild-type + empty vector), cWT+ATAD1 (wild-type expressing ATAD1) PEX3-/-(patient = empty vector), PEX3-/-+ATAD1 (PEX3-/-expressing ATAD1), cWT-ATAD1 (wildtype with ATAD1 deletion), PEX3-/-atad1-(patient with ATAD1 deletion). Statistical significance was calculated using the Welch’s test, p values are indicated as: ns (not significant), * p≤0.05, ** p≤0.01, *** p≤0.001, **** p≤0.0001. (E) Bar graph representation of calculated respiratory spare capacity (= uncoupled respiration minus basal respiration) of cWT (wild-type + empty vector), cWT+ATAD1 (wild-type expressing ATAD1) PEX3-/-(patient + empty vector), PEX3-/-+ATAD1 (PEX3-/-expressing ATAD1), cWT-ATAD1 (wildtype with ATAD1 deletion), PEX3-/-atad1-(patient with ATAD1 deletion). Statistical significance was calculated using the Welch’s test, p values are indicated as: ns (not significant), * p≤0.05, ** p≤0.01, *** p≤0.001, **** p≤0.0001. (F) Box-Whisker blot representation of the average (n=4) normalized peak intensity of all detected cardiolipin species, log10 pareto-scaled. (G) Box-Whisker blot representation of the average (n=4) normalized peak intensity of all detected phosphoethanolamin (PE) species, log10 pareto-scaled. (H) Box-Whisker blot representation of the average (n=4) normalized peak intensity of the subpopulation of phosphoethanolamin (PE) species causing the paired t-test significance in (G), log10 pareto-scaled. (I) Box-Whisker blot representation of the average (n=4) normalized peak intensity of all detected ether-phospholipid (plasmalogen) species, log10 pareto-scaled.

### Isolation of yeast mitochondria

Yeast cells were harvested at mid-log phase (OD600 = 2–3) unless indicated otherwise. Preparation of crude and purified mitochondria was described previously [51]. Briefly, the yeast pellet was washed once with MilliQ water, resuspended, and incubated in TD buffer (100 mM Tris-SO4, pH 9.4 and 100 mM DTT) for 15 min at 30°C. Spheroplasting was achieved by incubating cells in SP buffer (1.2 M sorbitol and 20 mM potassium phosphate, pH 7.4) supplemented with lyticase (2 mg/g of cell pellet) (Sigma-Aldrich) for 1 h 15 min at 30°C. Spheroplasts were gently washed in ice-cold SHE buffer (1.2 M sorbitol, 20 mM HEPES-KOH, pH 7.4, 2 mM MgCl2, 1 mM EGTA, and 1 mM PMSF) followed by homogenization in ice-cold SHE buffer (containing 0.6 M sorbitol) with a Dounce homogenizer (15-20 strokes). The crude mitochondrial fraction was obtained by differential centrifugation. Continuous Histodenz^™^ gradients were used to purify crude mitochondria. To make a gradient, 2.1 ml of 5, 10, 15, 20, and 25% Histodenz^™^ in SHE buffer was layered in 14 Å∼ 89 mm Ultra-Clear centrifuge tubes (Beckman) and the tubes were sat at 4°C for 3–4 h to allow the Histodenz^™^ to diffuse. Crude mitochondria were loaded on top of the chilled gradient and separated at 100,000 Å∼ g for 1 h at 4°C (SW41 rotor; Beckman). Intact purified mitochondria were recovered from a brown band at around 16% Histodenz^™^ concentration. Protein concentration was determined using a Bradford protein assay kit (Bio-Rad).

### Quantitative Proteomics

This method is developed in the lab of S.P.G.. Method description and summaries might therefore contain similar wording as published elsewhere.

### Sample preparation

Cell lysis and protein digestion, and peptide labeling: In order to fully use the sample multiplexing ability, an 10-plex experiment was designed containing the following proteome-wide comparisons: pex19Δ (n = 5), pex19Δmsp1Δ (n = 5), (we previously performed a 10-plex experiment with pex19Δ (n = 2), pex19Δmsp1Δ (n = 2) pex3Δ (n = 3), pex3Δmsp1Δ (n = 3) which is shown in the supplement (S-Fig.2). Cell pellets from each condition were resuspended at 4°C in buffer containing 8M urea, 50mM EPPS pH 8.5, 50mM NaCl, and protease inhibitors. Chilled zirconium oxide beads were added to cell slurries. Cells were lysed using a Mini-Beadbeater (Biospec products, Bartlesville, OK) at 4°C in 2mL screw cap tubes for 5 cycles of 30 s each, with 1 min pauses between cycles to avoid overheating. After centrifugation, clarified lysates were transferred to new tubes. Bicinchoninic acid (BCA) protein assay (Thermo Fisher Scientific) was performed to determine protein concentration. Proteins were then subjected to disulfide reduction with 5mM tris (2 carboxyethyl) phosphine (TCEP), (room temperature, 30 min) and alkylation with 10 mM iodoacetamide (room temperature, 30 min in the dark). 15 mM dithiotreitol was used to quench excess iodoacetamide (room temperature, 15 min in the dark). Proteins (200μg) were then chloroform/methanol precipitated and washed with methanol prior to air drying. Samples were resuspended in 8 M urea, 50 mM EPPS, pH 8.5, and then diluted to < 1M urea with 50mM EPPS, pH 8.5. Proteins were digested for 16 hours with LysC (1:100 enzyme:protein ratio) at room temperature, followed by trypsin (1:100 enzyme:protein ratio) for 6 hours at 37°C. Peptides were quantified using Pierce Quantitative Colorimetric Peptide Assay. TMT-10 reagents (0.8mg) were dissolved in 40μL anhydrous acetonitrile, and 7μL was used to label 70μg peptides in 30% (v/v) acetonitrile. Labeling proceeded for 1 hour at room temperature, until reaction was quenched using 7μL 5% hydroxylamine. TMT-labeled peptides were then pooled, vacuum centrifuged to dryness, and cleaned using 50mg Sep-Pak (Waters).

Offline basic pH reversed-phase (BPRP) fractionation: The pooled TMT-labeled peptide sample was fractionated using BPRP HPLC. We used an Agilent 1260 Infinity pump equipped with a degasser and a single wavelength detector (set at 220 nm). Peptides were subjected to a 50 min linear gradient from 8% to 40% acetonitrile in 10 mM ammonium bicarbonate pH 8 at a flow rate of 0.6 mL/min over an Agilent 300Extend C18 column (3.5 μm particles, 4.6 mm ID and 250 mm in length). We collected a total of 96 fractions, then consolidated those into 24 and vacuum centrifuged to dryness. Twelve of the 24 fractions were resuspended in a 5% acetonitrile, 1% formic acid solution. Fractions were desalted via StageTip, dried via vacuum centrifugation, and reconstituted in 5% acetonitrile, 5% formic acid for LC-MS/MS processing.

### Mass spectrometry for quantitative proteomics

Liquid chromatography and tandem mass spectrometry: Mass spectrometry data were collected using an Orbitrap Fusion Lumos mass spectrometer (Thermo Fisher Scientific) equipped with a Proxeon EASY-nLC 1000 liquid chromatography (LC) system (Thermo Fisher Scientific). Peptides were separated on a 100 μm inner diameter microcapillary column packed with ∼35 cm of Accucore C18 resin (2.6 μm, 150 Å, Thermo Fisher Scientific). Approximately 2μg peptides were separated using a 2.5 h gradient of acidic acetonitrile. We used the multinotch MS3-based TMT method (McAlister et al., 2014). The scan sequence began with a MS1 spectrum (Orbitrap analysis; resolution 120,000; mass range 400−1400 Th). MS2 analysis followed collision-induced dissociation (CID, CE = 35) with a maximum ion injection time of 150 ms and an isolation window of 0.5 Da. The 10 most abundant MS1 ions of charge states 2-6 were selected for MS2/MS3 analysis. To obtain quantitative information, MS3 precursors were fragmented by high-energy collision-induced dissociation (HCD, CE = 55) and analyzed in the Orbitrap (resolution was 50,000 at 200 Th) with a maximum ion injection time of 150 ms and a charge state-dependent variable isolation window of 0.7 to 1.2 Da [52].

### Data Analysis

MS2 mass spectra were processed using a SEQUEST-based software pipeline [21], [53], [54], [55]. Spectra were converted to mzXML using a modified version of ReAdW.exe. Database searching used the yeast proteome downloaded from Uniprot (UniProt-Consortium, 2015) in both forward and reverse directions, along with common contaminating protein sequences. Searches were performed using a peptide mass tolerance of 20 ppm, and a fragment ion tolerance of 0.9 Da. These wide-mass-tolerance windows were chosen to maximize sensitivity in conjunction with SEQUEST searches and linear discriminant analysis [56], [53]. TMT tags on lysine residues and peptide N termini (+229.163 Da) and carbamidomethylation of cysteine residues (+57.021 Da) were set as static modifications, while oxidation of methionine residues (+15.995 Da) was set as a variable modification.

Peptide-spectrum matches (PSMs) were adjusted to a 1% false discovery rate (FDR) [57]. Linear discriminant analysis was used to filter PSMs, as described previously [53], while considering the following parameters: XCorr, ΔCn, missed cleavages, adjusted PPM, peptide length, fraction of ions matched, charge state, and precursor mass accuracy. PSMs were identified, quantified, and collapsed to a 1% peptide false discovery rate (FDR) and then collapsed further to a final protein-level FDR of 1%. PSMs were quantified from MS3 scans; those with poor quality, MS3 spectra with total TMT reporter signal-to-noise ratio that is < 200, or no MS3 spectra were excluded from quantitation. Protein quantitation was performed by summing the signal-to-noise values for all peptides for a given protein. Each TMT channel was summed across all quantified proteins and normalized to enforce equal protein loading. Each protein’s quantitative measurement was then scaled to 100.

Quantification and Statistical Analysis: For proteomics experiments, the signal-to-noise ratios for each of the six channels were scaled to represent the percent of total signal in the six channels. Student’s t tests were performed in Microsoft Excel (2-tailed). P-values were adjusted using the Benjamini-Hochberg procedure for the 4,716 proteins that were tested. Significance was determined based on an adjusted P value of < 0.01.

### Quantitative proteomics data analysis for plotting

Quantitative proteomics samples were geometric mean-normalized. Common samples between experiments were not combined. Data for heatmaps were z-score normalized across each protein and plots were generated using the xpressplot.heatmap() function from XPRESSplot [58], [59]) and further formatting was performed using Matplotlib [60]. Volcano plots were generated using the geometric-normalized datasets with no further normalization. Plots were generated using the xpressplot.volcano() function [58], [59] and further formatting was performed using Matplotlib [60]. For code and raw data please refer to the ‘Data and Code Availability’ section.

### Western blot analysis

For immunostaining, proteins were transferred to nitrocellulose membrane after SDS-PAGE and PVDF membrane after BN-PAGE. The membranes were treated with rabbit/mouse antibodies against the tag or the chosen loading control according to standard procedures followed by secondary antibodies with fluorophore labels of HRP. The blots were scanned using a Licor Clx or developed following standard procedures. Final figure assembly was done using Adobe photoshop, where contrast and brightness were adjusted using only linear scaling. Final assembly was done in Adobe Illustrator. For S-Fig.2 the loading control signals were acquired on another gel running the same preparation instead of subsequent blotting.

### BN-PAGE

BN-PAGE was in collaboration with Ilka Wittig. Required method descriptions and summaries below might therefore contain similar language as published elsewhere.

### BN-PAGE

Mitochondrial pellets (1 mg mitochondrial protein) were resuspended in 100 μl solubilization buffer (30 mm HEPES, 150 mm potassium acetate, 10% [v/v] glycerol, and 5% [w/v] digitonin; protocol according to [61]. BN-PAGE was performed as published previously by [23]. Briefly, separation is achieved along a linear gradient of 3% to 18% polyacrylamide equivalent to a molecular mass range of 50 to 3000 kDa. BN gels were either directly stained with colloidal Coomassie Blue [62] or transferred to PVDF membranes for western blotting.

### Mammalian cell lines and culture

The fibroblast cell line PBD400-T1 was derived from a patient with peroxisomal biogenesis deficiency (Zellweger Syndrome). The patient carried a single nucleotide insertion (c542insT) leading to a premature stop codon in the peroxin gene, PEX3. An independent fibroblast cell line was derived from a patient with Zellweger syndrome carrying a terminating mutation, R176ter, in PEX16, (GM06231 cells, Coriell Institute). Both cell lines were a gift from S. Steinberg (Johns Hopkins Hospital, Baltimore, MA) and are now available through the Coriell Institute. Cells were immortalized with pBABE-hygro-hTert (a gift from Bob Weinberg (Addgene plasmid#1773; http://n2t.net/addgene:1773; RRID:Addgene_1773)) and sustained growth and survival was taken as readout for successful integration. Patient cells were infected, as described below, with retrovirus (pQXCIZ; Clontech) encoding V5-PEX3 or an empty vector control, and subsequently with retrovirus (pQXCIP; Clontech) encoding FLAG-HA-ATAD1 or an empty vector control.

Human fibroblast cells lines were maintained in DMEM/F12 (Gibco) with 15% FBS (Sigma Aldrich). HEK293T (Cat#CRL-3216^™^, ATCC) cells were maintained in DMEM (Corning) with 10% FBS and 1% penicillin/streptomycin (Gibco) (which was omitted in cells plated for virus production). Cells were cultured at 37°C with 5% CO_2_. Retrovirus was produced in HEK293T cells co-transfected with the vector containing the gene of interest, pUMVC-Gag-pol (a gift from Bob Weinberg (Addgene plasmid #8449; http://n2t.net/addgene:8449; RRID:Addgene_8449)), and pCMV-VSVG (a gift from Bob Weinberg (Addgene plasmid#8454; http://n2t.net/addgene:8454; RRID: Addgene_8454)) in a ratio of 3:2:1 using 1 mg/ml polyethylenimine (Cat#23966-1, Polysciences, Inc.). Retrovirus was harvested from the medium 48 h post-transfection, filtered through a 0.45 μm polyethersulfone membrane, collected, and stored at 4°C. Polybrene (Cat# TR-1003 G, EMD Millipore) was added to a final concentration of 10 μg/mL and a 1:1 mixture of viral supernatant and medium (DMEM + 15% FBS) was applied to the target cells. 24 h post-infection, target cells were selected using 4 μg/ml puromycin (ThermoFisher) or 150 μg/ml zeocin (Invivogen) for 5–7 days. Stable cell lines were maintained in the appropriate medium with 0.5 μg/ml puromycin (ThermoFisher) and/or 20 μg/ml zeocin (Invivogen).

Lentivirus (for making PEX-13-GFP/ PEX10-GFP expressing lines) was produced in HEK293T cells that were co-transfected with the vector containing the gene of interest and second generation envelope and packaging plasmids (psPAX–gift from Didier Trono (Addgene plasmid#12260; http://n2t.net/addgene:12260; RRID:Addgene_12260) and pMD2.G–a gift from Didier Trono (Addgene plasmid#12259; http://n2t.net/addgene:12259; RRID:Addgene_12259) in a ratio of 4:2:1 using polyethylenimine (PEI) (Cat#23966-1, Polysciences, Inc.) (PEI:DNA mass ratio 3:1). Virus-containing medium was collected after 48 h, filtered through 0.45 mm polyethersulfone membrane, aliquoted, and stored at −80°C. Lentivirus-containing supernatant was added in a 1:1 ratio with medium (DMEM + 15% FBS) to cells together with polybrene transfection reagent at a final concentration 8-10 mg/mL (Cat# TR-1003-G, EMD Millipore). 48 h post-infection, GFP+ cells were sorted using the Aria Cell Sorter. Cells were allowed to recover for 1 to 3 days before imaging.

### CRISPR-Cas9 deletion of *ATAD1*

Oligonucleotides encoding sgRNAs targeting human *ATAD1* were annealed and cloned into the Px458 vector. pSpCas9(BB)-2A-GFP (PX458) was a gift from Feng Zhang (Addgene plasmid # 48138; http://n2t.net/addgene:48138; RRID:Addgene_48138). Two sgRNAs were used, targeting either exon 2 (sense oligo: 5’-**CACC**CCGACTCAAAGGACGAGAAA-3’) or exon 5 (sense oligo: 5’-**CACC**CGGTCAGTGTCGAAGGCTGA-3’; overhang in bold). Patient fibroblasts (see above) were transfected with one of these two plasmids using Lipofectamine 3000 (ThermoFisher). Two days later, GFP+ cells were sorted and plated as single cells into each well of 96 well plates. Clonal cell lines were maintained in DMEM + 15% FBS, expanded, and loss of ATAD1 was assessed by western blot using a monoclonal ATAD1 antibody (Cat# 75-157 Antibodies Incorporated/NeuroMab, Davis, CA, USA).

### Electron microscopy

Cells were grown to 70-80% confluency in a 6 well tissue culture dish, washed with Milli-Q water and fixed with formaldehyde solution. Staining was performed with 0.75% uranyl formate. The images were recorded on a 120 kV Tecnai G2 Spirit transmission electron microscope (FEI) using an Ultrascan 4000 4k × 4k CCD camera (Gatan) at 6,000× magnification at the specimen level. Approximately 50-100 mitochondria were collected from each specimen, and the experiments were repeated with three cultures in each case. Data were analyzed by an individuum via visual identification of mitochondrial morphologies (cWT n=232, PEX pex3-/atad1-n=253, PEX pex3-/atad1+ n=266). Their morphology appearance was described as mitochondria with many cristae and low electron density (type 1), mitochondria with fewer cristae (type 2) and damaged mitochondria with high electron density and/or almost no cristae (type 3) (representative images are displayed in Fig.4B and S-Fig.5). Representative images for figures 4B and 5 were assembled using GraphPad Prism6 and 8, respectively.

### Mitochondrial oxygen consumption rate (OCR)

Mitochondrial oxygen consumption rate (OCR) was assessed using the generated cell lines in an XF96e Extracellular Flux Analyzer (Seahorse Bioscience), as described previously [63]. Culture media of human fibroblast cells plated with a cell count of 1000 cells per well in an XF96 cell culture microplates were replaced with XF Dulbecco’s modified Eagle medium (DMEM) containing 10 mM glucose, 2 mM L-glutamine (Life Technologies), and 2 mM sodium pyruvate (Life Technologies). OCR was measured at 37°C with 1-min mix, and 3-min measurement protocol. OCR was analyzed after 30 min incubation in a CO_2_-free incubator. Oligomycin (2 uM), carbonilcyanide m-fluorophenylhydrazone (FCCP) (1.5uM), and rotenone + antimycin (1uM each) were sequentially injected into each well to assess basal respiration, coupling of respiratory chain, and mitochondrial respiratory capacity. OCRs were normalized relative to cell count in each well. Basal respiration and maximal respiration were calculated by subtraction of the non-mitochondrial respiration from the initial respiration and respiration after FCCP uncoupling, respectively. The respiratory spare capacity was calculated by substraction of the basal respiration from the maximal respiration. All blots are generated from raw XFe96e Flux analyzer data from 2 independent experiments with 4 technical replicates on each day (n=8) in WAVE software and processed through the XF Mito Stress Test Report. Significance was calculated with the Welch’s-test in GraphPad Prism8 and blots were created in GraphPad Prism8.

### Lipidomics

This method description is widely distributed at the University of Utah and external collaborators. The following method statements/ summaries given by the University of Utah metabolomics core might therefore contain similar language as published elsewhere.

### Lipidomics Sample Extraction

Samples were prepared as described in [64] with following modifications: Solutions are pre-chilled on ice. Lipids are extracted in a solution of 225 µL MeOH and 750 µL methyl tert-butyl ether (MTBE) containing internal standards (Avanti SPLASH LipidoMix, Avanti LM6003 Cardiolipin Mix and Avanti Ceramide LipidoMix all at 10 µL per sample; LPC(26:0)-d4 at 156 pmol per sample; C18(plasm)-18:1(d9) PE at 135 pmol per sample; C18(plasm)-18:1(d9) PC at 128 pmol per sample). For other than occasional vortexing samples incubate on ice for 1 h. Following the addition of 188 µL of PBS and brief vortexing the samples rest at room temperature for 10 min. Samples are centrifuged at 16,000 x *g* for 5 minutes at 4 °C to collect the upper phases which then evaporated to dryness under speedvac. The bottom aqueous layer is collected separately and dried under speedvac. Lipid extracts are reconstituted in 500 µL IPA and transferred to an LC-MS vial for analysis. Concurrently, a process blank sample and pooled quality control (QC) samples are prepared by taking equal volumes (∼50µL) from each sample after final resuspension.

### Mass Spectrometry Analysis of Samples

Lipid extracts are separated on a Waters Acuity UPLC CSH C18 1.7 µm 2.1 × 100 mm column maintained at 65 °C connected to an Agilent HiP 1290 Sampler, Agilent 1290 Infinity pump, equipped with an Agilent 1290 Flex Cube and Agilent 6530 Accurate Mass Q-TOF dual AJS-ESI mass spectrometer. For positive mode, the source gas temperature is set to 225 °C, with a drying gas flow of 11 L/min, nebulizer pressure of 40 psig, sheath gas temp of 350 °C and sheath gas flow of 11 L/min. VCap voltage is set at 3500 V, nozzle voltage 1000V, fragmentor at 110 V, skimmer at 85 V and octopole RF peak at 750 V. For negative mode, the source gas temperature is set to 300 °C, with a drying gas flow of 11 L/min, a nebulizer pressure of 30 psig, sheath gas temp of 350 °C and sheath gas flow 11 L/min. VCap voltage is set at 3500 V, nozzle voltage 2000V, fragmentor at 100 V, skimmer at 65 V and octopole RF peak at 750 V. Samples are analyzed in a randomized order in both positive and negative ionization modes in separate experiments acquiring with the scan range m/z 100 – 1700. Mobile phase A consists of ACN:H_2_O (60:40 *v/v*) in 10 mM ammonium formate and 0.1% formic acid, and mobile phase B consists of IPA:ACN:H_2_O (90:9:1 *v/v*) in 10 mM ammonium formate and 0.1% formic acid. The chromatography gradient for both positive and negative modes starts at 15% mobile phase B then increases to 30% B over 2.4 min, it then increases to 48% B from 2.4-3.0 min, then increases to 82% B from 3 – 13.2 min, then increases to 99% B from 13.2-13.8 min where it’s held until 16.7 min and then returned to the initial conditioned and equilibrated for 5 min. Flow is 0.4 mL/min throughout, injection volume is 3 µL for positive and 10 µL negative mode. Tandem mass spectrometry is conducted using the same LC gradient at collision energy of 25 V.

### Analysis of Mass Spectrometry Data

QC samples (n=8) and blanks (n=4) are injected throughout the sample queue and ensure the reliability of acquired lipidomics data. Results from LC-MS experiments are collected using Agilent Mass Hunter (MH) Workstation and analyzed using the software packages MH Qual, MH Quant, and Lipid Annotator (Agilent Technologies, Inc.). Results from the positive and negative ionization modes from Lipid Annotator are merged then split based on the class of lipid identified. The data table exported from MHQuant is evaluated using Excel where initial lipid targets are parsed based on the following criteria. Only lipids with relative standard deviations (RSD) less than 30% in QC samples are used for data analysis. Additionally, only lipids with background AUC counts in process blanks that are less than 30% of QC are used for data analysis. The parsed excel data tables are normalized to tissue mass and positive and negative mode data are merged.

### Data Pretreatment

Data was analyzed using in-house software to normalize and scale (pareto). Supplemental tables also contain the Raw Input in which no treatment of any kind was performed.

### Resource Availability

All unique/stable reagents and resources generated within this study are available from the corresponding author, Jared Rutter, upon request and without restriction.

### Data and Code Availability

The quantitative proteomics data have been deposited to the ProteomeXchange Consortium via the PRIDE (Perez-Riverol et al., 2019) partner repository with the dataset identifier {pending}. The code and processed data tables used to analyze these quantitative proteomics data and reproduce the associated plots are available at GitHub https://github.com/j-berg/nuebel_2020. RNA-Seq and ribosome-footprint profiling data have been deposited at the Gene Expression Omnibus (http://www.ncbi.nlm.nih.gov/geo) under accession number GSE{pending}, and will be made publicly available before publication.

## Material and methods

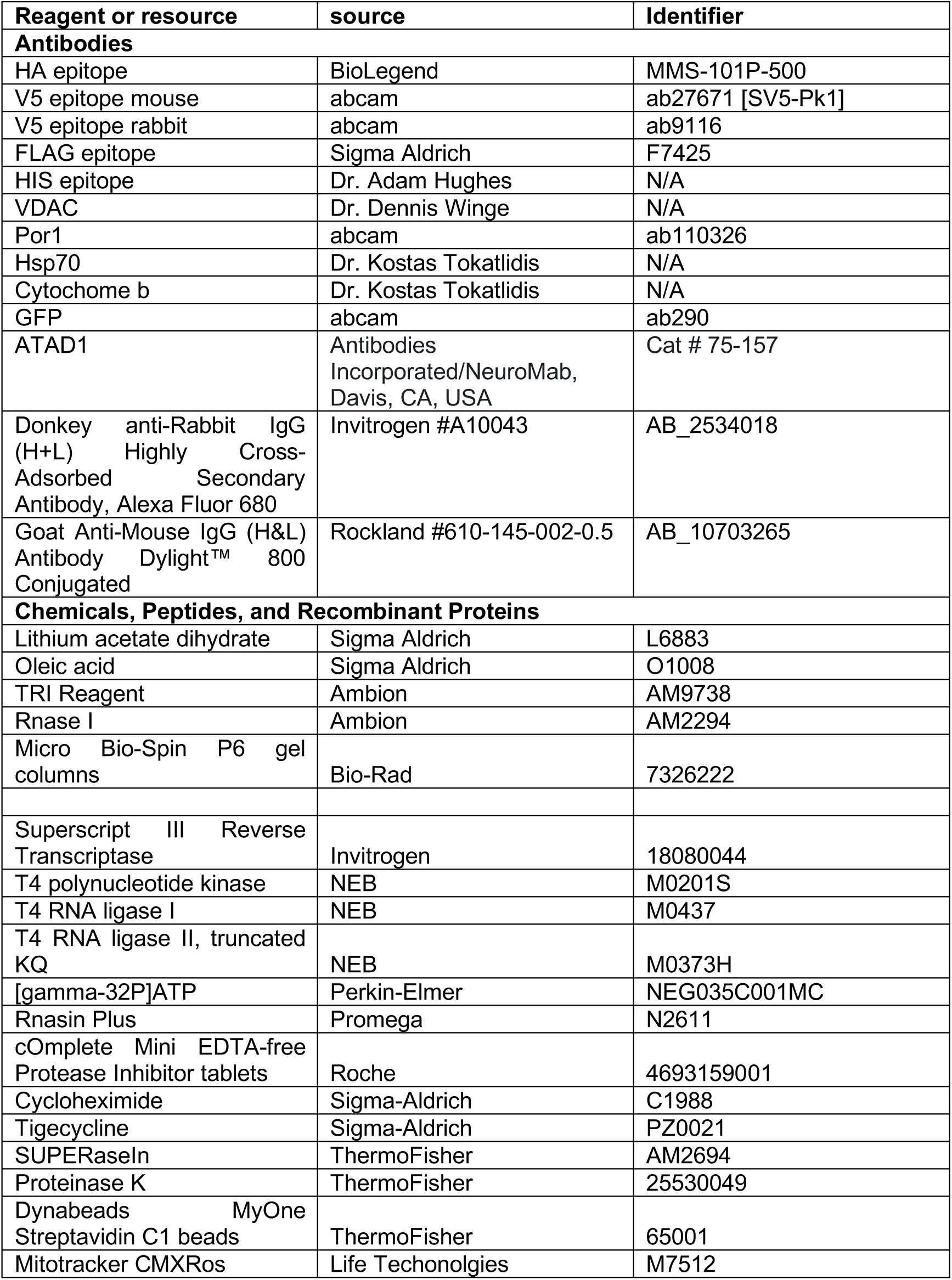
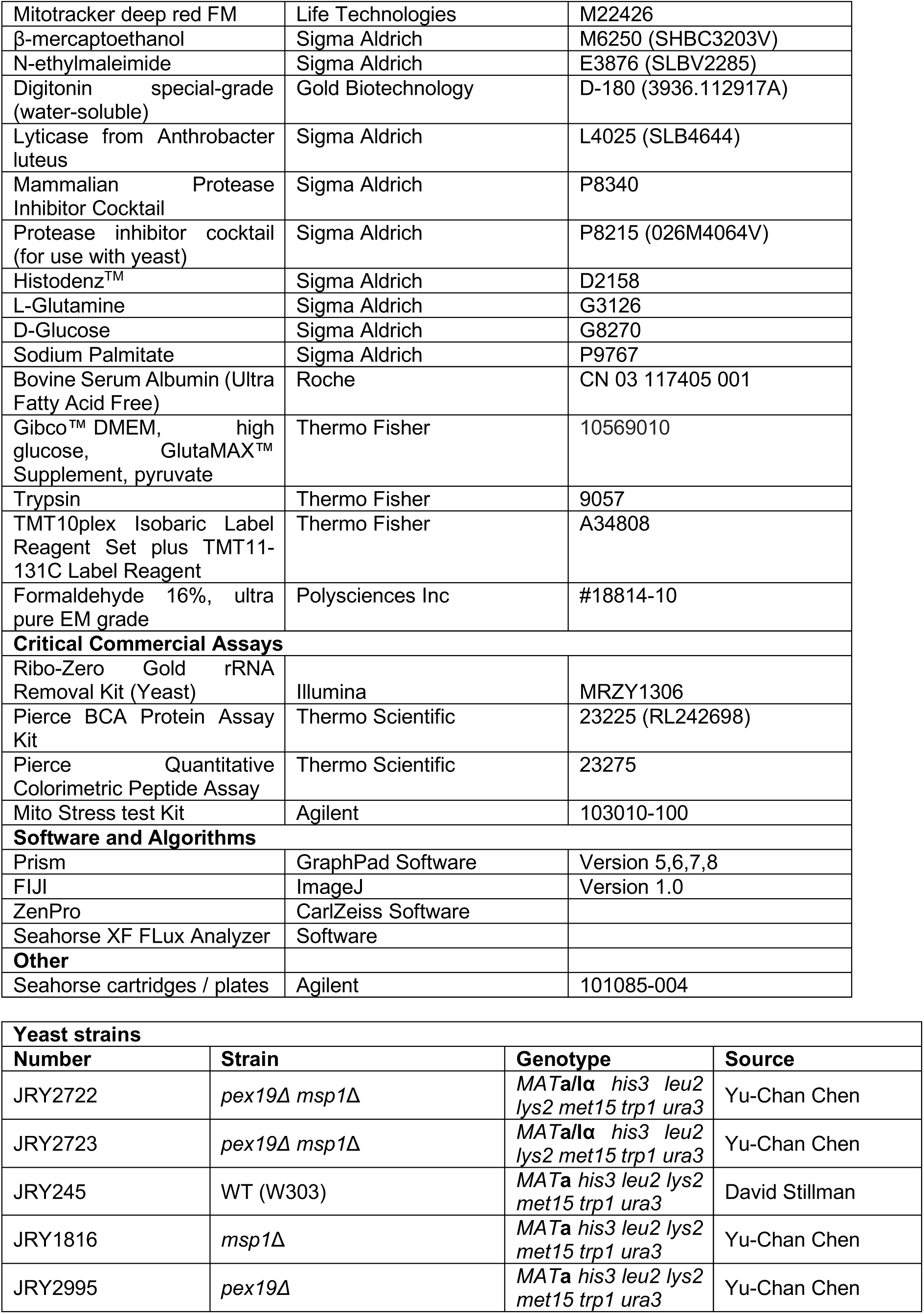
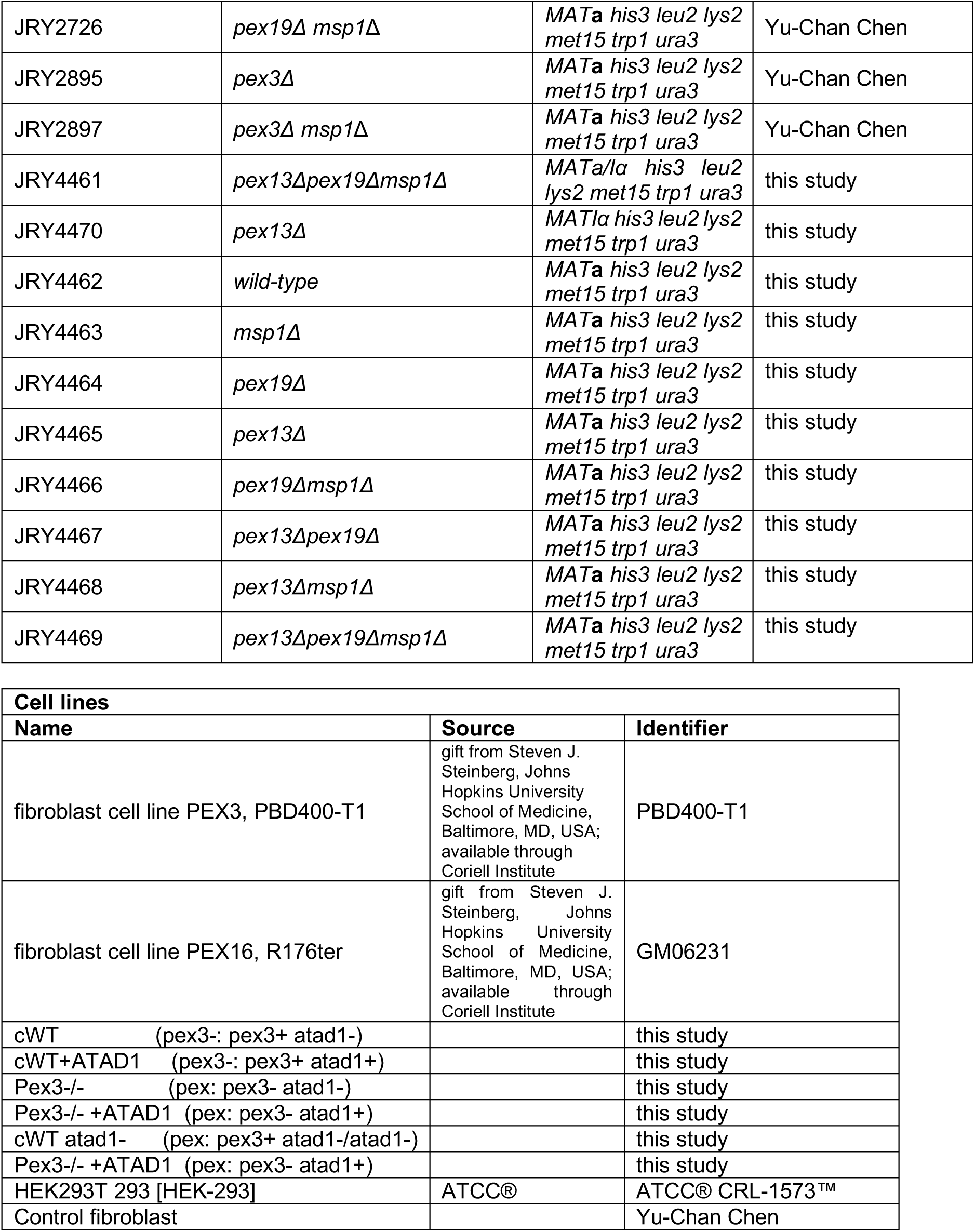

## Abbreviations

PBDs: Peroxisomal Biogenesis Disorders
ZSD: Zellweger Spectrum Disease
ATP: Adenosine-tri-phosphate
RNA-Seq: RNA-Sequencing
TE: translational efficiency
GET: gated entry of tail anchored proteins
SDS: sodium-dodecyl-sulfate
BN: blue native
PAGE: polyacryl-gel-electrophoresis
EM: electron microscopy
ER: endoplasmatic reticulum
PerMit: peroxisomal and mitochondrial tether

## Acknowledgements

This work was supported by the Howard Hughes Medical Institute (to J.R.), the NIH grant 1RO1GM115174-01A (to J.R.), a grant from the Nora Eccles Treadwell Foundation (to J.R.), the Global Foundation for Peroxisomal Disorders and the Wynne Mateffi Foundation (to J.R.), the Deutsche Forschungsgemeinschaft: SFB 815/Z1 (to I. W.) and the BMBF mitoNET -German Network for Mitochondrial Disorders 01GM1113B (to I.W.). This study was supported by the University of Utah’s Electron Microscopy Facility, Metabolomics Facility, Metabolism Profiling Facility, Flow Cytometry Facility and the High-Throughput Genomics Shared Resource at the Huntsman Cancer Institute. We thank Jana Meisterknecht for technical assistance regarding native electrophoresis, Linda Nikolova for technical assistance in performing electron microscopy data, John Alan Maschnek and James Cox for lipidomics, Anil Laxman for metabolism profiling support, James Marvin for assistance at the flow core, and Katja Dove for critically reading the manuscript. Some antibodies were generously gifted by Dr. Kostas Tokatlidis.

J.A.B. received support from NIDDK T32DK11096601 to Wendy W. Chapman and Simon J. Fisher and is supported by the National Cancer Institute of the National Institutes of Health under Award Number F99CA253744. J.E.C. is funded by S10OD016232, S10OD021505, and U54DK110858.

S.P.G. is funded by GM97645. N.B. and C.A. are funded by the Canadian Institutes of Health Research 34575 (to N.B.) and the Richard and Edith Strauss Foundation (to C.A.). The content of this manuscript is solely the responsibility of the authors and might not represent the official views of the National Institutes of Health.

J.R., E.N. and Y.C.C submitted an international patent application regarding the mechanism described in this paper. The authors declare no further competing interests.

## Author contributions

E.N. and J.R., designed the study. E.N. wrote the first version of the manuscript. J.R. and J.M. contributed significantly to finish the written manuscript. E.N., S.F., J.T.M., and Y.C.C. collected the data. E.N. conceived of the presented idea, developed the theory and carried out most of the experiments. Y.C.C. created yeast strains and generated plasmid constructs, and imaged human cell lines. S.F. generated and confirmed plasmid constructs, yeast strains and generated the human cell lines used in this manuscript and imaged yeast cells. J.T.M. performed the ribosome profiling and assembled corresponding figures. S.L. produced lentivirus and infected cells, K.J.C. performed and analyzed quantitative mass spectrometry under the mentorship of S.P.G. I.W. performed and analyzed the Blue Native PAGE. J.M.W. generated and confirmed the human ATAD1ko cell lines. C.U.K. performed imaging on the Airyscan (Zeiss) while E.N. prepared cell staining for imaging. J.A.B. performed data analysis of the quantitative proteomics data and prepared figures for these data. N. B., C. A. and L.C. advised on peroxisomal biochemistry and the clinical impact of presented data. All authors commented and approved of the final manuscript.

---

**Supplemental Figure to Figure 1 (S-Fig. 1).**
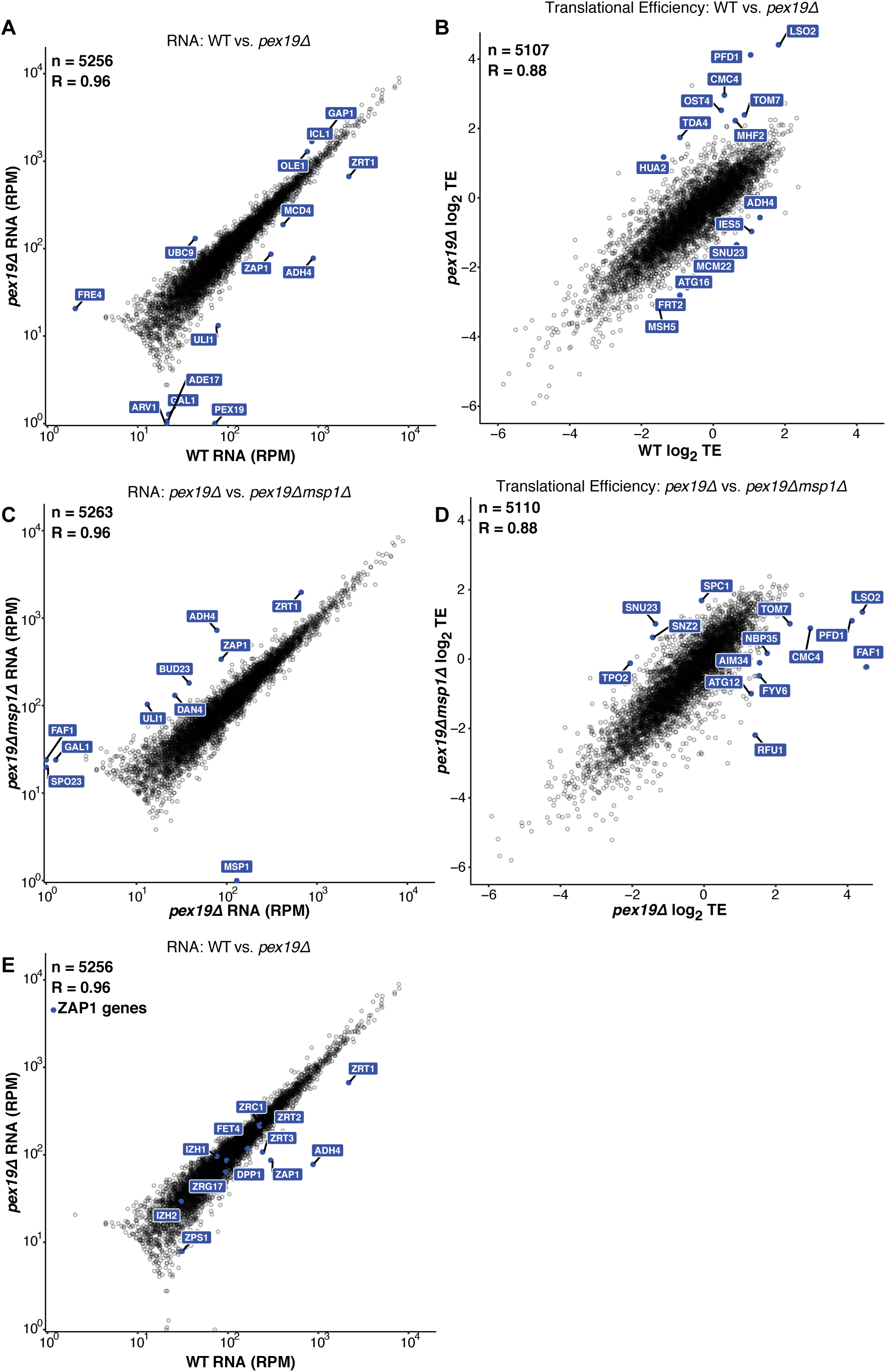
(A) Wild-type versus *pex19Δ* RNA levels are blotted. All outliers are indicated. (B) Translational efficiency (TE) of wild-type versus *pex19Δ*. All outliers are indicated. (C) *pex19Δ* versus *pex19Δmsp1Δ* RNA levels are blotted. All outliers are indicated. (D) Translational efficiency (TE) of *pex19Δ* versus *pex19Δmsp1Δ*. All outliers are indicated. (E) ZAP1 target genes abundance at the RNA level in wild-type versus pex19Δ.

**Supplemental Figure to Figure 2 (S-Fig. 2).**
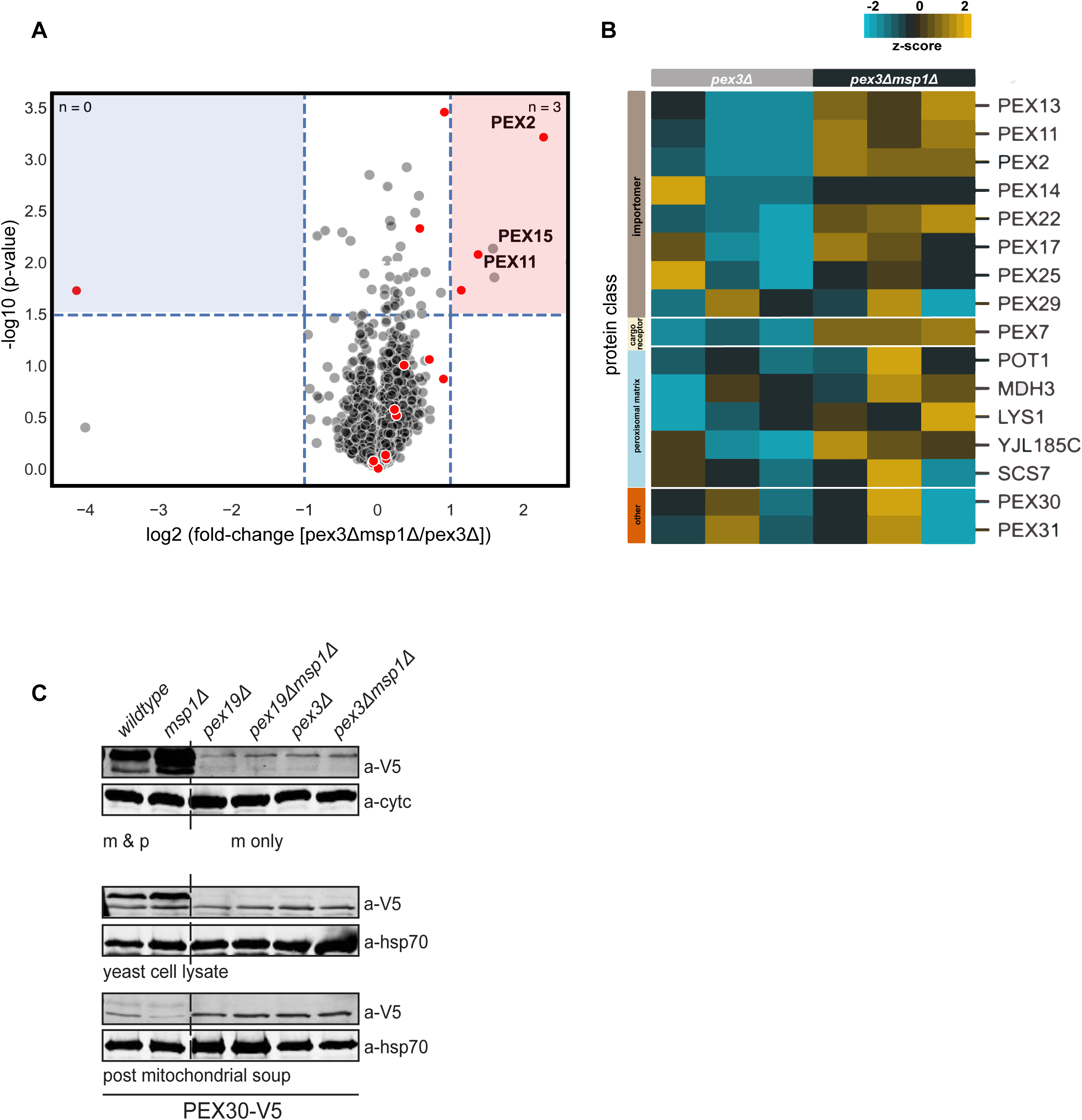
(A) Volcano plot (log^10^) representing the average of 3 biological replicates of each strain (pex3Δ and pex3Δmsp1Δ) to indicate most enriched/ decreased proteins in the mitochondrial proteome of pex3Δ and pex3Δmsp1Δ. Detected by quantitative mass spectrometry. P-values were corrected using the Benjamini-Hochberg adjustment procedure. Gray dots represent all measured proteins, and red dots highlight peroxisomal-associated proteins. (E) Heatmap (log^2^) representing 3 biological replicates of each strain (pex3Δ and pex3Δmsp1Δ) and their peroxin protein levels detected by quantitative mass spectrometry, protein classes are indicated in the row labels. Values were z-score normalized by protein. (F) Total yeast cell lysate, post mitochondrial soup and Histodenz^™^ purified mitochondria of each strain (pex19Δ and pex19Δmsp1Δ; pex3Δ pex3Δmsp1Δ) transformed with Pex30^V5^ (expressed by its endogenous promotor) were separated by SDS-PAGE and immunoblotted for Pex30 (a-V5) cytochrome c (cyt c) and HSP70 (hsp70) were used as loading control.

**Supplemental Figure to Figure 3 (S-Fig.3).**
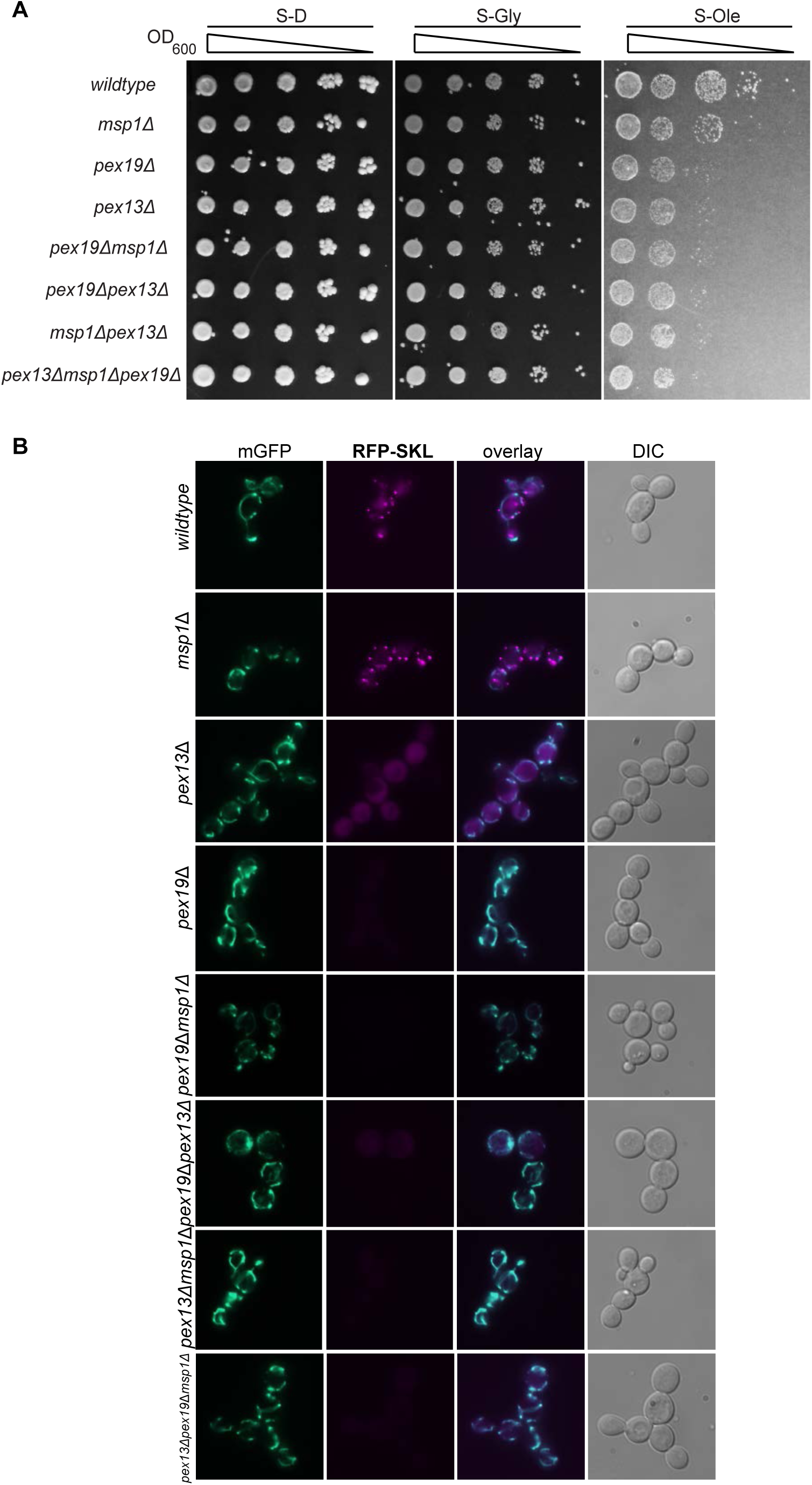
(A) Yeast cells of each strain (*wild-type, msp1Δ, pex19Δ, pex13Δ pex19Δmsp1Δ, pex13Δmsp1Δ, pex13Δpex19Δ and pex13Δpex19Δ msp1)* were grown to mid-log phase, back diluted to 1 OD and serial dilutions were dropped onto agar plates containing synthetic media with dextrose (S-D), glycerol (S-Gly), and oleate (S-Ole). (B) Yeast cells of each strain (*wild-type, msp1Δ, pex19Δ, pex13Δ pex19Δmsp1Δ, pex13Δmsp1Δ, pex13Δpex19Δ and pex13Δpex19Δ msp1)* expressing RFP-SKL (peroxisomal marker) construct and mitochondria-targeted green fluorescent protein (GFP) were grown to mid-log phase and analyzed by fluorescence microscopy. Representative images are shown. DIC, differential interference contrast.

**Supplemental Figure to Figure 4 (S-Fig.4).**
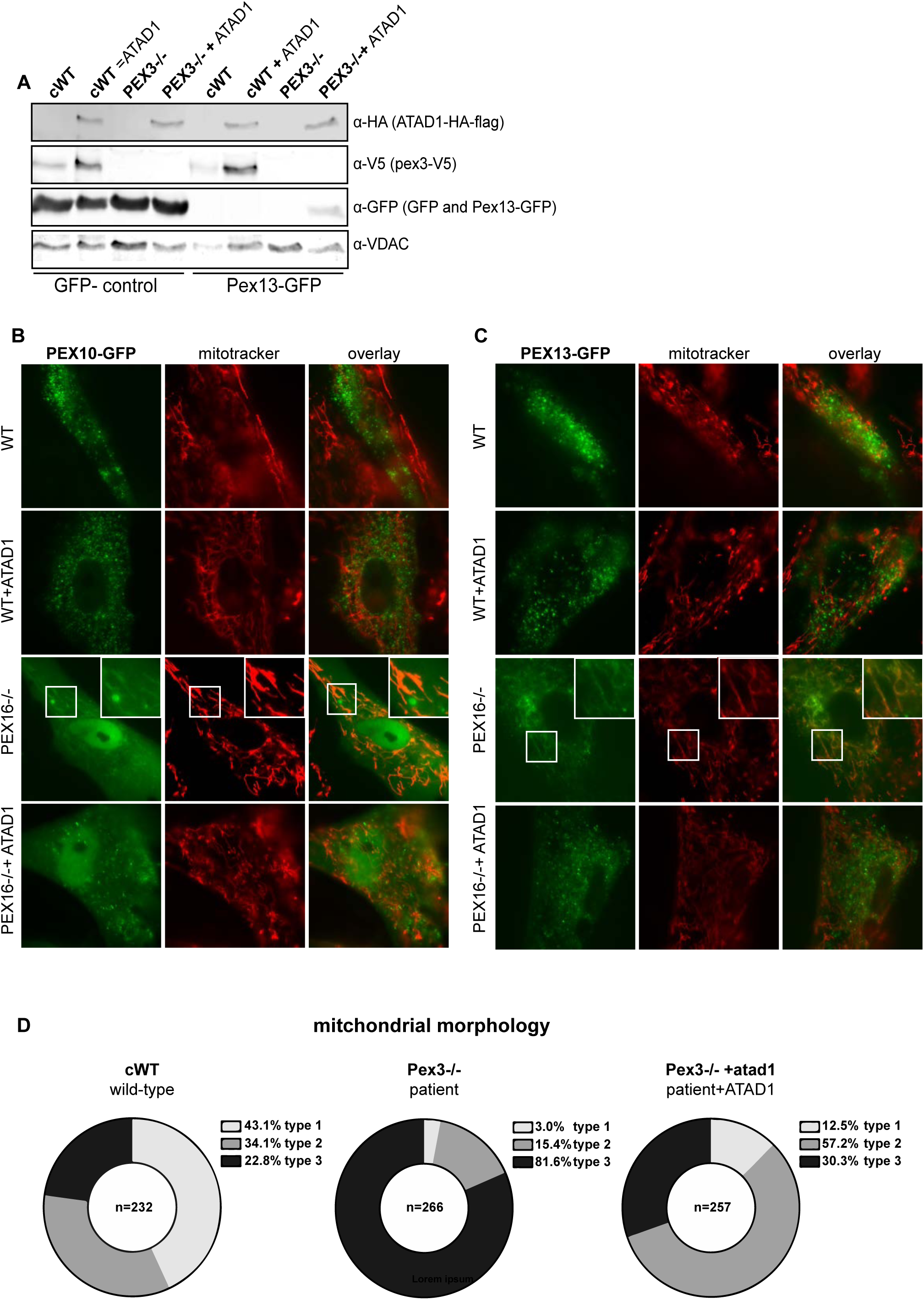
(A) 25 ug human cell lysate of cWT (wild-type + empty vector), cWT+ATAD1 (wild-type expressing ATAD1) PEX3-/-(patient + empty vector), PEX3-/-+ATAD1 (PEX3-expressing ATAD1), cWT-ATAD1 (wildtype with ATAD1 deletion), PEX3-/-atad1-(patient with ATAD1 deletion) transfected with control^GFP^ or Pex13^GFP^ were separated by 10% SDS-PAGE and immunoblotted for ATAD1-HA-flag with a-HA, Pex3^V5^ with a-V5, and for GFP with a-GFP. VDAC was visualized with a-VDAC as loading control. (B) Live fluorescence microscopy of patient fibroblast cell lines: WT (wild-type + empty vector), WT+ATAD1 (wild-type expressing ATAD1) PEX16-/-(patient + empty vector), PEX16-/-+ATAD1 (PEX16-/-expressing ATAD1) expressing PEX10-GFP stained with Mitotracker red (MT) to visualize the mitochondrial network and GFP fused to PEX10 (PEX10-GFP) to investigate the localization. Images are reprinted from [26] with permission of Yu-Chan Chen. (C) Live fluorescence microscopy of patient fibroblast cell lines: WT (wild-type + empty vector), WT+ATAD1 (wild-type expressing ATAD1) PEX16-/-(patient = empty vector), PEX16-/-+ATAD1 (PEX16-/-expressing ATAD1) expressing PEX13-GFP stained with Mitotracker red (MT) to visualize the mitochondrial network and GFP fused to PEX13 (PEX13-GFP) to investigate the localization. Images are reprinted from [26] with permission of Yu-Chan Chen. (C) Quantification of electron microscopy imaging. 3 independent experiments (different passage number and independent fixing and staining procedure) of cWT (wild-type + empty vector), PEX3-/-(patient + empty vector), and PEX3-/-+ATAD1 (PEX3-/-expressing ATAD1) were quantified. Mitochondria were counted (cWT (wild-type + empty vector) n=232, PEX3-(patient + empty vector) n=266, and PEX3-/-+ATAD1 (PEX3-/-expressing ATAD1) n=257) and binned into 3 types (mitochondria with many cristae and low electron density (type 1), mitochondria with fewer cristae (type 2) and damaged mitochondria with high electron density and/or almost no cristae (type 3)) representing different morphology phenotypes. The results are displayed in % values.

**Supplemental Figure to EM microscopy methods section (S-Fig. 5).**
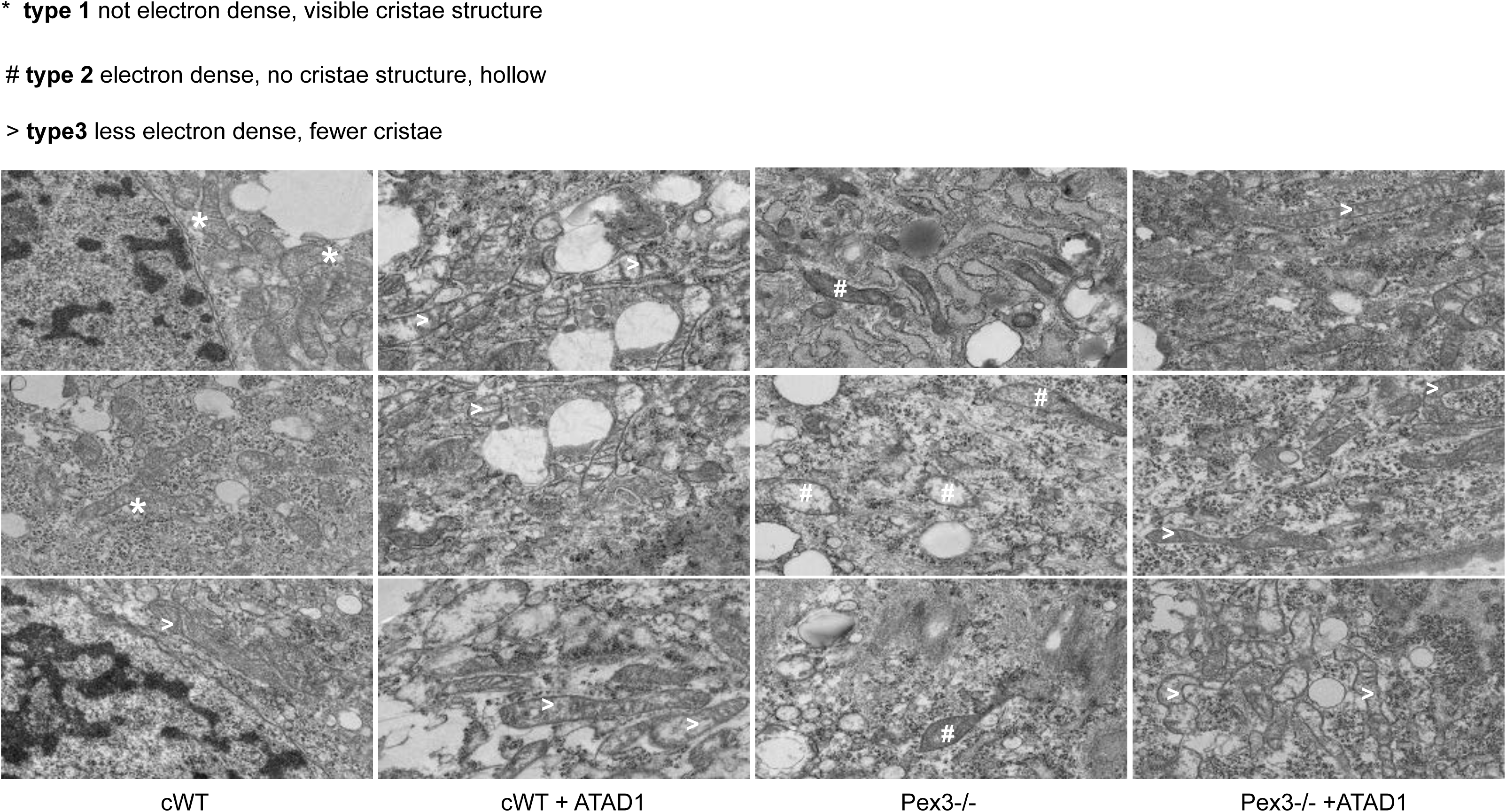
Electron microscopy of patient fibroblast cell lines: cWT (wild-type + empty vector), cWT+ATAD1 (wild-type expressing ATAD1) PEX3-/-(patient + empty vector), PEX3-/-+ATAD1 (PEX3-expressing ATAD1). Representative images of the different observed mitochondrial morphologies are shown.

**Supplementary Table 1 for Figure 4 (S-Table 1)**

Sheet 1 (Raw CL) Excel compilation of raw lipidomics values for cardiplipin species

Sheet 2 (Raw PE) Excel compilation of raw lipidomics values for PE species

Sheet 3 (Raw ether-phospho-lipids) Excel compilation of raw lipidomics values for ether-phospho-lipid species

Sheet 4 (normalized and pareto scaled CL) Excel compilation of normalized and scaled lipidomics values for cardiolipin species

Sheet 5 (normalized and pareto scaled PE) Excel compilation of normalized and scaled lipidomics values for PE species

Sheet 6 (normalized and pareto scaled CL) Excel compilation of normalized and scaled lipidomics values for ether-phospho-lipid species

## Paper highlights

- The absence of peroxisomes maintains peroxin levels on transcriptional and translational level.
- Several peroxins (Pex13, Pex11, Pex14, Pex2, Pex17, Pex25), accumulate on the mitochondrial outer membrane and reveal the formation of the peroxisomal docking complex.
- To our knowledge this is the first evidence of a membrane protein translocase capable of relocation to another organelle and thus putatively interfering with its function.
- ATAD1 is sufficient to restore mitochondrial morphology, respiration, and metabolism in cells derived from PBD patients.
- This work will redirect the focus of PBD therapies towards ameliorating mitochondrial dysfunction to help lower symptom burden.

